# Maturation of HIV-1 neutralizing antibodies in a germinal center conditional expression mouse model

**DOI:** 10.64898/2026.03.30.715358

**Authors:** Ming Tian, Jillian Davis, Hwei-Ling Cheng, Lily M. Thompson, Marie-Elen Tuchel, Aimee Chapdelaine Williams, Audrey Yin, Bailey Wilder, Ivy DiBiase, Michael S. Seaman, Frederick W. Alt

## Abstract

In germinal centers, activated B cells modify their antigen receptors through somatic hypermutation (SHM), followed by antigenic selection that favors expansion of high affinity B cells. The affinity maturation process is critical for development of broadly neutralizing antibodies (bnAbs) against the human immunodeficiency virus-1 (HIV-1). BnAbs have been isolated from some people living with HIV-1. Because these antibodies target conserved epitopes of the HIV-1 Envelope (Env) protein, they inhibit a broad spectrum of viruses. Eliciting bnAbs by vaccination is a top priority for HIV-1 prevention, but reproducing the lengthy maturation of bnAbs is a major challenge. The problem is typified by VRC01 class antibodies, which recognize the CD4 binding site of HIV-1 Env protein. To reach the CD4 binding site, antibodies need to navigate through adjacent glycans. Accommodating the glycans requires multiple SHMs in germinal center (GC) B cells, including infrequent events. For this reason, VRC01 vaccine development often stalls at this point. We have generated a mouse model aimed at providing a potential solution for navigating this vaccine design impediment. To this end, we made a mouse model that expresses a stalled VRC01 intermediate conditionally in GC B cells. This system has three advantages: 1) direct expression of the intermediate obviates prior immunization steps, thereby shortening the immunization scheme; 2) the conditional expression system bypasses tolerance control checkpoints that sometimes delete B cells expressing bnAbs; 3) the intermediate responds to immunization in GCs, the physiological site of affinity maturation. With this model, we established an immunization method to mature the VRC01 intermediate into heterologous neutralizing antibodies against viruses with a native glycan shield. Since high mutation load is common among bnAbs, the germinal center conditional expression system could provide a general tool for boost immunogen design to overcome roadblocks in the maturation pathway.

**Author summary:** In response to antigenic stimulation, cognate B cells become activated and form germinal centers in lymphoid tissues. Germinal center B cells modify their antigen receptors through somatic hypermutation (SHM) of immunoglobulin variable region gene exons, with antigen selecting for high affinity B cells by providing survival advantage. This mechanism accounts for antibody affinity maturaton over the gradual course of an immune response. Affinity maturation is critical for generating potent, neutralizing antibodies against diverse strains of the human immunodeficiency virus-1 (HIV-1). These broadly neutralizing antibodies (bnAbs) are heavily mutated, reflecting lengthy affinity maturation over years of chronic infection. Recapitulating the affinity maturation process is a major challenge for bnAb induction by vaccination. In immunization experiments, bnAb development often stalls at rate limiting steps that involve infrequent, but functionally important, mutational events. Overcoming such obstacles requires boost immunogens that can stimulate the stalled B cells to acquire the requisite mutations. To this end, we recapitulated the maturation arrest of a bnAb lineage by expressing a stalled antibody in mouse germinal center B cells. Using this mouse model, we developed boost immunization conditions that advanced the antibody maturation beyond a roadblock to attain neutralizing activities against heterogenous viruses.

## Introduction

Naïve B cells express B cell receptors (BCRs) that are encoded by germline immunoglobulin (Ig) variable region exon gene segments assembled by V(D)J recombination. Such germline BCRs usually have low antigen binding affinities. After B cell activation, Ig variable regions accumulate somatic hypermutations (SHMs) in germinal center (GC) B cells (1). The GC microenvironment selects for mutations that improve antigen binding affinity. The combination of SHM and antigenic selection gradually evolves the germline precursor into mature antibodies with high antigen binding affinities. Affinity maturation is especially important for broadly neutralizing antibodies (bnAbs) against the human immunodeficiency virus-1 (HIV-1) (2). These antibodies can inhibit diverse HIV-1 strains by recognizing conserved epitopes of Envelope (Env) protein. BnAbs have been found in some individuals living with HIV-1. A major goal of HIV-1 vaccine development is to induce similar antibodies. However, it has been difficult to reproduce the extensive affinity maturation involved in bnAb evolution, especially the steps that require infrequent SHM events, such as insertions and deletions (“indels”) or mutation of sequences that are poor targets (“cold spots”) of the SHM proccess. Acquisition of such improbable, but functionally important, mutations is rate-limiting for bnAb development (3). Since SHM is not evenly targeted across variable region exon sequences, it is unfeasible to direct SHMs to desired positions in immunoglobulin (Ig) variable region exons. Overcoming maturation hurdles requires immunogens that can boost SHM activity and select for requisite mutations. Boost immunogens target GC or memory B cells that express intermediate BCRs between the germline precursor and mature bnAbs. Intermediates represent a continuum of antibodies with different degrees of affinity maturation. Boost immunogen design needs to focus on specific antibodies. In this regard, the intermediates that are stalled at rate limiting steps are pertinent targets.

The efficacy of boost immunogens is critically dependent on their affinity for target BCRs. On the one hand, the immunogen should bind to target BCR with high enough affinity to drive SHM in GCs. On the other hand, if the affinity is too high, the immunogen will not exert sufficient selective pressure for further maturation. The optimal affinity can only be determined empirically by immunization, and mouse models are widely used for this purpose. Many mouse models have been made to express germline precursors of various bnAb lineages, but these mouse models are not ideal for testing boost immunogens for specific intermediates. Starting from precursor, it may take several immunization steps to reach the target intermediate. These steps not only take time but also increase variability. The progression of an immune response is usually asynchronous in different animals. When a boost immunogen is administered at a given point, it will encounter heterogeneous intermediates, which may have a range of affinities for the boost immunogen. It will be hard to correlate immunization outcome with antigen-binding affinity. Both issues, time and variability, become more serious for late intermediates. One way to address these issues is to express the target intermediate for a boost immunogen directly in a mouse model.

The conventional strategy to express a specific antibody in a mouse model is to integrate its heavy chain (HC) and light chain (LC) variable region coding sequences into the J_H_ and Jκ loci respectively. This strategy has two potential caveats for expressing bnAb intermediates. First, the intermediate will be expressed constitutively in B cells, starting from the progenitor stage. There have been multiple precedents where B cells expressing bnAb precursors were deleted by developmental checkpoints (4–7). The problem is likely due to negative selection against poly- or auto-reactivities, which appear to be more commonly associated with bnAbs than other antibodies (8). Since bnAb intermediates normally arise *de novo* in GC B cells, they should not be subject to central tolerance control, and deletion of intermediate B cells in bone marrow is a nonphysiological event. B cell developmental defects may not be associated with all bnAb intermediates, and even under negative selection, some B cells may still survive to maturity. However, such mouse models have a second issue: intermediate BCRs will respond to boost immunogens in the context of naive B cell. The antigenic response of naive B cells may differ from those of GC or memory B cells, which are the physiological target of boost immunogens (9–11). Besides intrinsic differences, GC and naïve B cells have distinct competitors. Normally, bnAb intermediate B cells compete with antigenically selected fellow GC B cells. When bnAb intermediates are expressed in naïve B cells, their competitors are diverse naïve B cells. The T cell environment is also different between the two scenarios. Boost immunization can benefit from existing T cell help in an established GC, whereas immunization of naïve mice needs to also prime T cell response *de novo*.

To address the concerns discussed above, we devised a method to express bnAb intermediates conditionally in GC B cells in a mouse model. The GC conditional expression strategy allows expression of intermediates to skip progenitor and naive B cell stages, thereby avoiding tolerance control checkpoints. More important, boost immunogens will engage intermediate BCRs in the physiological context of GC. As a test of the method, we generated such a mouse model to mature an intermediate of the VRC01 class antibodies, which target the CD4 binding site of Env protein (12, 13). Substantial progress has been made in eliciting VRC01 class antibodies in mouse models (14–26) and, more recently, in human clinical trials (27–29). A main obstacle in VRC01 development is to accommodate glycans that restrict access to the CD4 binding site. Overcoming the glycan barrier remains a challenge for boost immunogen design. To solve this problem, we incorporated a VRC01 intermediate stalled at a rate-limiting step into the GC conditional expression mouse model. With this mouse model, we established boost immunization methods that matured the VRC01 intermediate into heterologous neutralizing antibodies against natively glycosylated viruses.

## Results

### GC conditional expression of bnAb intermediate in mouse model

The GC conditional mouse model contained conditional expression cassettes for the heavy chain (HC) and light chain (LC) alleles of a bnAb intermediate. The HC cassette contained two pre-rearranged variable region exons ((VDJ)_H_, Fig 1A) in tandem. Each variable region was preceded by a V_H_ promoter that normally transcribes mouse variable regions. Based on their relative distance to the downstream μ constant region (Cμ), the proximal and distal variable regions corresponded to the germline (GL) precursor and the intermediate antibody (IA) of a bnAb HC, respectively. Two polyadenylation sites (pAx2, Fig 1A) were interposed between the distal and proximal (VDJ)_H_s. The polyadenylation sites can truncate the distal IA (VDJ)_H_ transcript. Furthermore, based on relative distance, the associated intronic enhancer (iEμ, Fig 1A) should preferentially activate the proximal V_H_ promoter for the GL-(VDJ)_H_. Thus, in the default configuration of the cassette, the proximal GL-(VDJ)_H_ would be expressed in progenitor and naïve B cells. To switch on the expression of the distal IA-(VDJ)_H_ in GC B cells, we flanked the proximal GL (VDJ)_H_ and the polyadenylation sites with two loxP sites (Fig 1A). In combination with a GC-specific cre transgene, the floxed region can be deleted conditionally in GC B cells to express the distal IA (VDJ)_H_. The strategy used to conditionally express the IA-(VDJ)_H_ in GC B cells was also applied to conditionally express the LC cassette as well (see below).

**Fig 1.**
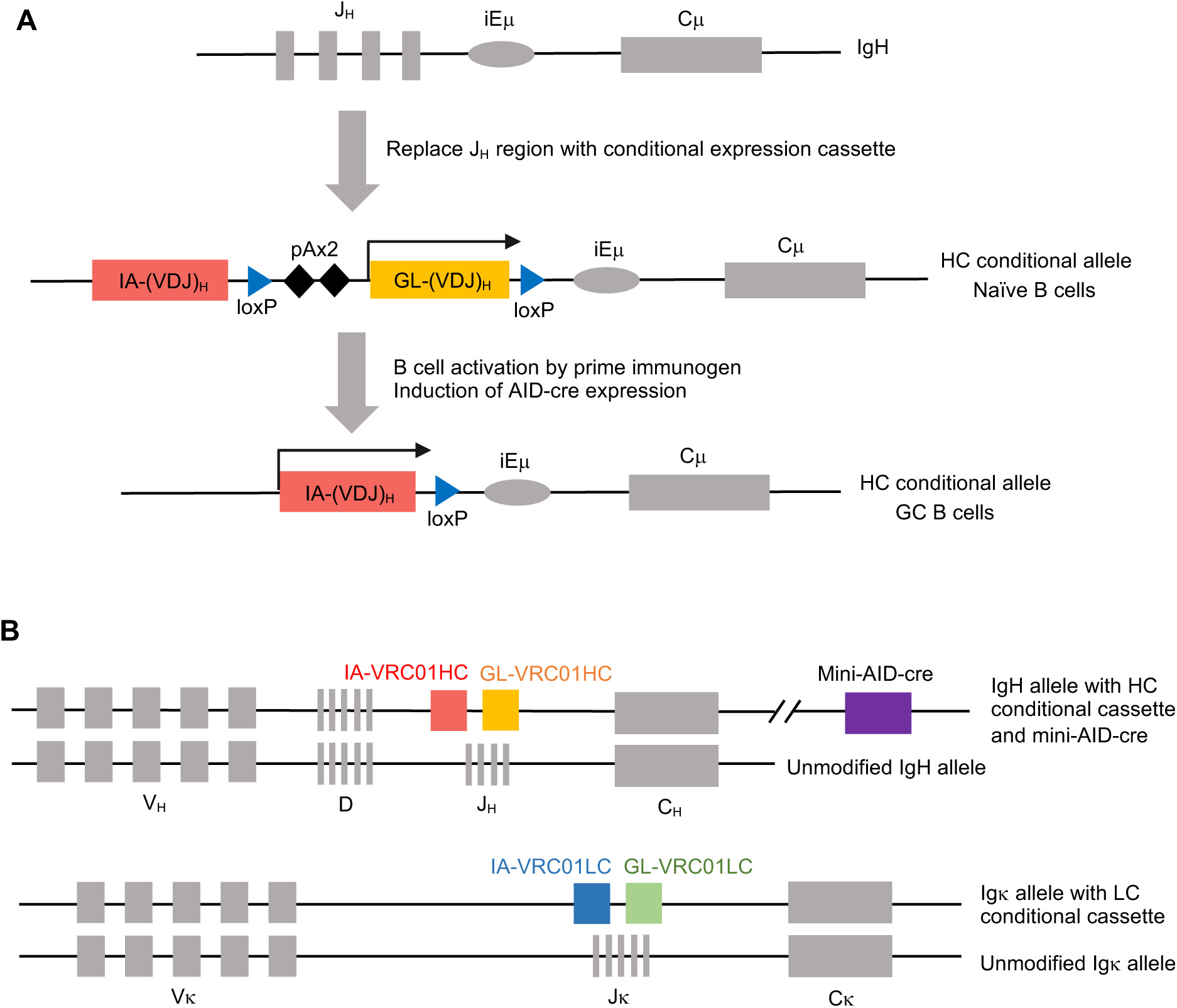
Diagram of the GC conditional expression cassette. (A) Mechanism of GC conditional expression of bnAb intermediate antibody HC. GL-(VDJ)_H_ and IA-(VDJ)_H_ represent the coding exons for germline and intermediate variable regions of a bnAb. Transcription of GL or IA-(VDJ)_H_ is indicated by arrow. pAx2 represents two tandem polyadenylation signals. iEμ represents the intronic enhancer of IgH. (B) Components of the GC conditional expression mouse model for a VRC01 intermediate. GL- and IA-VRC01HC represent the coding exons for the variable regions of GL- and IA-VRC01HC respectively. GL-and IA-VRC01LC represent the coding exons for the variable regions of GL- and IA-VRC01LC respectively. Mini-AID-cre transgene was integrated downstream of the IgH locus.

AID-cre (30, 31), S1pr2-cre (32) and GCET-TamCre (33) transgenes have been used for conditional deletion of floxed region in GC B cells. In these transgenes, the cre cDNA is either integrated into the endogenous loci (*Aicda* (30) or *Gcet* (33)) or embedded in bacterial artificial chromosomes (BACs) of *Aicda* (31) or *S1pr2* loci (32). In such contexts, cre expression is under the control of *Aicda*, *S1pr2* or *Gcet* transcriptional elements, which are active in GC B cells. Although these cre transgenic mouse lines are available, the GC-cre transgene resides on a different chromosome from the HC and LC cassettes, and it would take extra breeding steps to combine all three components. To simplify the breeding scheme, we integrated a GC-cre transgene into the same IgH allele as the HC cassette. Since the existing large BAC-based cre transgene is difficult for site-specific integration, we generated a streamlined version of AID-cre transgene by concatenating defined *Aicda* transcriptional elements (34, 35) and cre cDNA (S1 Table). We integrated the mini-AID-cre transgene via homologous recombination into a site downstream of a cluster of CTCF binding elements (CBEs), which mark the 3’ end of IgH locus (36). The 3’ CBEs can insulate the mini-AID cre transgene from the transcriptional enhancers of the IgH loci (37) so that cre expression would follow the pattern of AID rather than IgH. To test the function of the mini-AID-cre, we combined the transgene with a YFP reporter, the expression of which is dependent on excision of a STOP cassette by cre recombinase (38) (S1A Fig). We analyzed YFP expression in the mouse that contained both the mini-AID-cre transgene and the YFP reporter. We found that, in the unimmunized mouse, YFP expression was off in naive B cells (B220^+^IgD^+^, S1B Fig). In response to immunization with sheep red blood cells, which stimulated strong GC formation, about 30% GC B cells (B220^+^CD38^lo^GL7^+^CD95^+^) were YFP^+^. These results showed that the mini-AID-cre transgene can mediate GC-specific deletion of floxed cassette, and its activity was comparable to that of the cre transgene based on *Aicda* BAC (31).

Having validated the mini-AID-cre transgene, we used the system to conditionally express a VRC01 intermediate in GC B cells. The 1538-79 intermediate antibody is the most advanced VRC01 class antibody from an earlier immunization experiment in a VRC01 mouse model (16). As mentioned in the Introduction, glycans near the CD4 binding site are barriers to VRC01 antibodies, and the N-linked glycan at Env residue 276 (N276 glycan) is particularly obstructive. The 1538-79 antibody can only neutralize virus that lacks the N276 glycan. Similar antibodies have been elicited in other immunization experiments or during HIV-1 infection (16, 20–22, 24–26, 39–43). Thus, the 1538-79 antibody can serve as a representative of a VRC01 intermediate that is stalled at this roadblock. Hereafter, we refer to the 1538-79 antibody as IA-VRC01. The HC and LC of the IA-VRC01 contain 8.4% and 2.9% mutations respectively at the DNA level. Reversion of the mutations gave rise to germline-reverted VRC01 (GL-VRC01). We incorporated the VDJ exons of the GL-VRC01 and IA-VRC01 into the proximal and distal positions of the conditional expression cassette for HC and LC (Fig 1B, S1 Table). We integrated the HC cassette into the J_H_ locus of the IgH allele that contained the mini-AID-cre transgene. As explained above, linkage of the HC cassette and mini-AID-cre on the same chromosome can simplify mouse breeding to reconstitute the complete mouse model. We generated a separate mouse line with the LC cassette at the Jκ locus. By breeding the HC/mini-AID-cre and LC mouse lines together, we reconstituted the complete mouse model (Fig 1B), which is referred to as the GC model hereafter.

We used flow cytometry to analyze the splenic B cells of the GC model. The spleen of the GC model had similar proportions of B and T cells as unmodified wild-type (WT) mice (Fig 2A, 2B and 2F). However, the GC model did exhibit some noticeable differences in B cell subtypes from wild-type mice (Fig 2A, 2C-2F). These differences are likely attributable to the expression of pre-rearranged GL-VRC01 in most B cells of the GC model. The largely monoclonal B cells in the GC model expressed more uniform levels of surface IgM (Fig 2C) and Igκ (Fig 2D) relative to the polyclonal B cells in WT mice. Because of isotype exclusion (44), expression of VRC01LC as Igκ diminished Igλ B cells (Fig 2D and 2F). Compared to polyclonal BCRs in WT mice, the GL-VRC01 BCR also directed more B cells into the marginal zone B cell compartment, but follicular B cells remained dominant in the GC model (Fig 2E and 2F). Overall, the GC model had largely normal splenic B cell populations.

**Fig 2.**
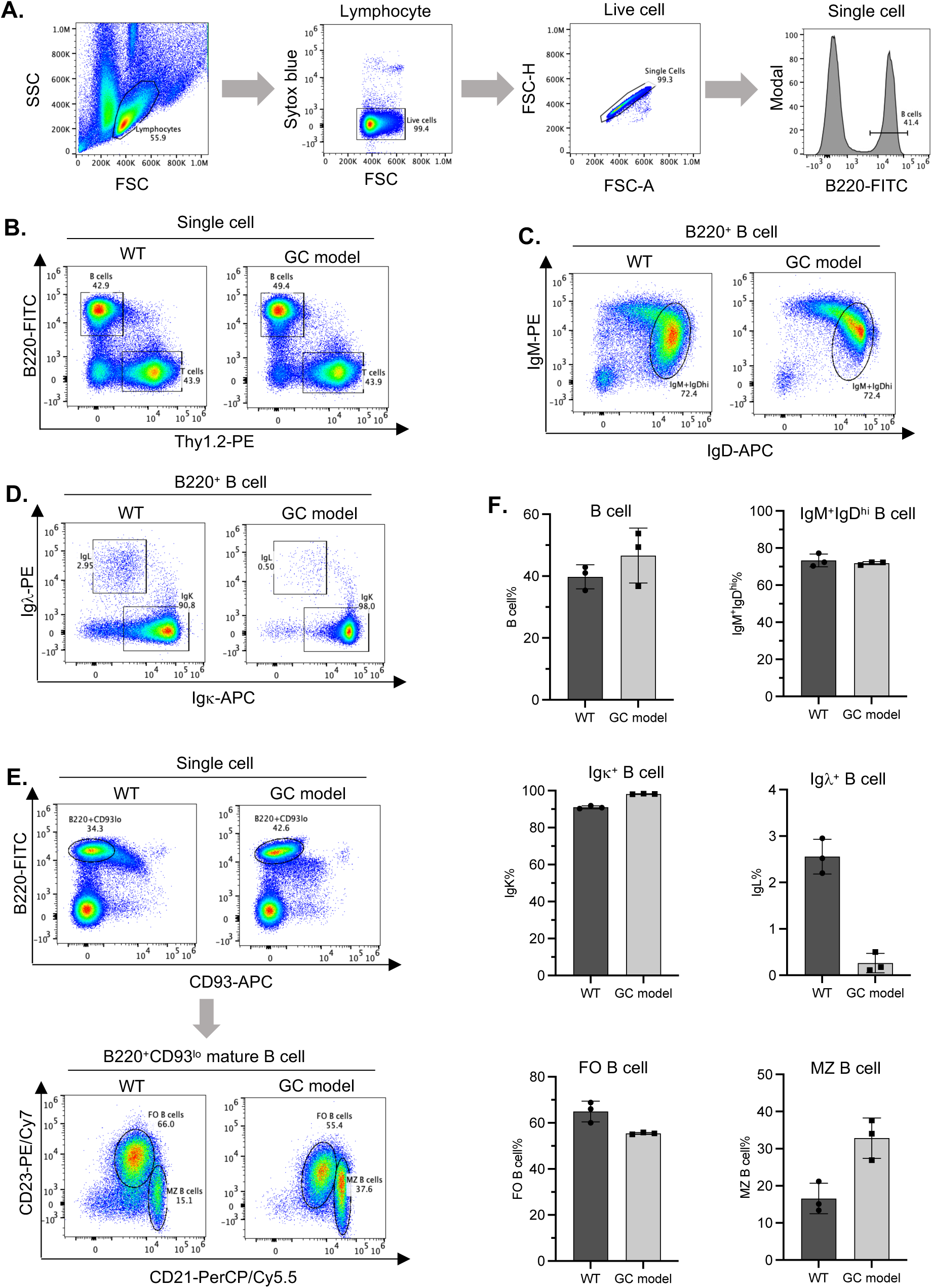
Flow cytometric analysis of splenic B cells from the GC conditional model. (A) FACS plots and gating strategy preceding the analysis in panels B-E. Analysis of splenocytes from a WT mouse is shown here as an example. The gate from the preceding plot is indicated above the plots. Gating scheme: lymphocyte>live cells>single cells>B220^+^ B cells. (B) Total B cells and T cells. Gating scheme: lymphocyte>live cells>single cells>B220^+^Thy1.2^-^(B cells); B220^-^Thy1.2^+^ (T cells). (C) Surface IgM/IgD expression. Gating scheme: lymphocyte>live cells>single cells>B220^+^ B cells>IgM^+^IgD^hi^ (naive B cells). (D) Igκ^+^ and Igλ^+^ B cells. Gating scheme: lymphocyte>live cells>single cells>B220^+^ B cells>Igκ^+^Igλ^-^. (E) Follicular (FO) and marginal zone (MZ) B cells. Gating scheme: lymphocyte>live cells>single cells>B220^+^CD93^lo^ mature B cells>CD21^lo^CD23^hi^ (follicular B cell); CD21^hi^CD23^lo^ (marginal zone B cell). **F**. Summary of the flow cytometric analysis. The bar graphs summarize the flow cytometric analysis, as shown in panels B-E, of 3 WT mice and 3 GC model mice. The bar height corresponds to the mean of the 3 samples, and the error bar represents standard deviation.

Based on design, naive B cells of the GC model should express GL-VRC01. For validation, we used eOD-GT8 (14) as a probe to detect VRC01 BCR expression. eOD-GT8 is an engineered form of Env gp120 subunit that binds strongly to VRC01 class precursors. We found that more than 90% of splenic B cells in the GC model stained positive for the eOD-GT8 probe (Fig 3A and 3B), and this interaction was abrogated by mutations in the CD4 binding site (ΔeOD-GT8, Fig 3A). However, eOD-GT8 staining cannot differentiate GL-VRC01 from IA-VRC01 expression because it bound equally to both, based on Enzyme-Linked Immunosorbent Assay (ELISA, Fig 3D). GL-VRC01 or IA-VRC01 expression could be ascertained definitively at the sequence level. To this end, we sorted single naive splenic B cells (B220^+^IgM^+^IgD^hi^) from the GC model (Fig 3C), performed single-cell RT-PCR to amplify VRC01HC and LC cDNAs from the sorted B cells, and used both Sanger sequencing (S2 Fig, S2 Table) and restriction digest (S3 Fig) to distinguish GL-VRC01 from IA-VRC01 transcript. Based on analysis of 238 B cells from 5 mice, all of the B cells expressed GL-VRC01HC (Fig 3C and S4 Fig). In one mouse, 10/49 B cells expressed IA-VRC01LC (S4A Fig), and the remaining B cells in this and the other 4 mice expressed GL-VRC01LC (Fig 3C and S4A Fig). The incidence of IA-VRC01LC expression likely resulted from leaky expression of AID-cre transgene in naïve B cells. In these cases, B cells expressed BCRs that were composed of GL-VRC01HC and IA-VRC01LC. Based on ELISA, the binding activity of the hybrid antibody, GL-VRC01HC/IA-VRC01LC, behaved identically to GL-VRC01, whereas the reciprocal hybrid antibody, IA-VRC01HC/GL-VRC01LC was equivalent to IA-VRC01 (S4C Fig). Thus, somatic hypermutations in IA-VRC01HC accounts for affinity maturation of this antibody. Leaky expression of IA-VRC01 in some naïve B cells would not generate BCRs that behave like IA-VRC01.

**Fig 3.**
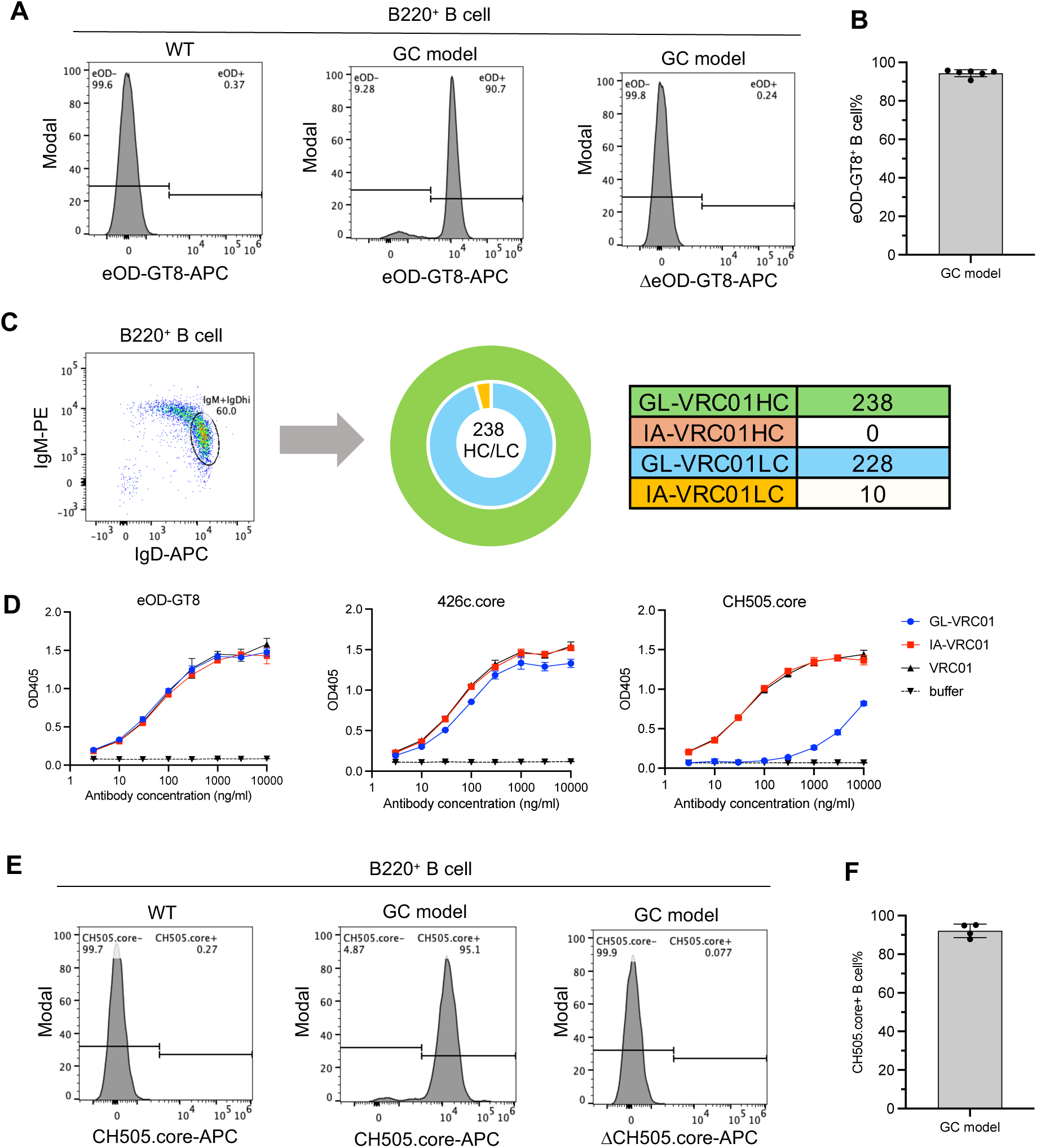
Expression of GL-VRC01 in naïve B cells of the GC model. (A) Flow cytometric analysis of eOD-GT8 binding to splenic B cells. Gating scheme: lymphocyte>live cells>single cells>B220^+^ B cells >eOD-GT8^+^, ΔeOD-GT8^-^ (VRC01 class B cells). (B) The percentage of eOD-GT8 binding splenic B cells. The bar graph summaries the results of flow cytometric analysis of 6 mice of the GC model, as shown in panel A (middle FACS plot). Each dot represents one mouse. The height of the bar corresponds to the mean, and the error bar represents standard deviation. (C) sc-RT-PCR analysis of naive B cells from the GC model. The FACS plot shows the naïve B cell population (B220^+^IgM^+^IgD^hi^) that was sorted for sc-RT-PCR. The pie chart summarizes the proportion of HC (outer ring) and LC (inner ring) that corresponded to GL-VRC01 or IA-VRC01. In total, 238 HC/LC pairs from five mice were analyzed. Color codes are shown in the adjacent table, which also lists the number of HC and LC in each category. The shade of the color over the number correlates with HC or LC frequency: 100%, full color; 0%, no color. The identifies of the HC and LC were determined by Sanger sequencing (S2 Fig) or restriction digest (S3 Fig) of the PCR product. The full data are in S4A Fig. (D) ELISA measurement of the interactions of GL-VRC01, IA-VRC01 and mature VRC01 antibodies with eOD-GT8, 426c.core and CH505.core. The plots show ELISA titration curves, which are color coded for GL-VRC01, IA-VRC01 and VRC01, as indicated to the right; buffer served as the baseline. Coating antigen is indicated at the top of each plot. The curve corresponds to the average of three technical replicates, and the error bar represents standard deviation. (E and F) Flow cytometric analysis of CH505.core binding to splenic B cells of the GC model. The plots are labeled in the same way as for panels A and B.

### Induction of IA-VRC01 expression in GC B cells by prime immunization

Induction of IA-VRC01 expression in GC B cells required a prime immunogen that bound to GL-VRC01 with sufficient affinity to activate GL-VRC01 naïve B cells. Furthermore, the prime immunogen should bind more strongly to IA-VRC01 than GL-VRC01. The affinity differential would favor the expansion of B cells that have switched from GL-VRC01 to IA-VRC01 expression in GC. Like other GC-specific cre transgenes (30–33), the mini-AID-cre transgene mediates partial deletion of floxed cassette in GC B cells (S1C Fig). By giving the IA-VRC01 B cells selective advantage, the immunogen could increase the proportion of IA-VRC01 B cells in GCs. In addition to affinity, antigen multimerization can strengthen BCR interaction through avidity effects. Many platforms are available for antigen multimerization. Based on our prior experience, we chose antibody Fc region fusion to dimerize fusion partners (45). All of the antigens in the experiments below were in the form of Fc fusions.

eOD-GT8 is an effective prime immunogen for VRC01 precursor B cells (14, 46), but its affinities for GL-VRC01 and IA-VRC01 were equal (Fig 3D, left plot). 426c.core gp120 is another prime immunogen for VRC01 precursors (18). The 426c immunogen is derived from the Env gp120 subunit of a clade C virus, 426c. Conversion of 426c.gp120 to 426c.core involves mutation of 3 glycosylation sites near the CD4 binding site and truncation of V1-V3 loops. These modifications can open access to the CD4 binding site (47, 48). Like eOD-GT8, 426c.core also interacted strongly with both GL-VRC01 and IA-VRC01 (Fig 3, center plot). The IA-VRC01 was originally elicited by 426c-based immunogens (16). The GL-VRC01 retained the CDR3 of the IA-VRC01. Since the CDR3 of the IA-VRC01 may have undergone affinity maturation, it probably contributed to the interaction of GL-VRC01 with the 426c.core. Based on this reasoning, we predicted that heterologous gp120 might bind differentially to GL-VRC01 versus IA-VRC01. To test this prediction, we generated gp120.cores from several viral isolates; these heterologous cores had analogous glycosylation site mutations and V1-V3 loop truncations as the 426c.core. We tested the binding of these gp120.cores to GL-VRC01 and IA-VRC01 by ELISA and found that the gp120.core of a clade C virus, CH505 transmission founder (CH505TF) (49), had the desired properties for prime immunization. CH505.core associated more strongly with IA-VRC01 than GL-VRC01 (Fig 3D, right panel). Despite its weak interaction with GL-VRC01, it stained similar proportions of B cells as eOD-GT8 in the GC model, and the staining was specific to the CD4 binding site (Fig 3E and 3F). Therefore, we chose CH505.core as the prime immunogen to activate GL-VRC01 B cells in the GC model.

The abundance of GL-VRC01 B cells (>90%) in the GC model presented a non-physiological condition for immunization. A solution for this problem was to adoptively transfer B cells from the GC model to WT mice. This method is widely used to titrate bnAb precursor B cell frequency in recipient mice (50, 51). The GC model was derived from a F1 ES cell line, which is of mixed 129/Sv and C57BL/6 background. To avoid graft rejection, we used F1 progeny of 129/Sv and C57BL/6 mice as recipients for B cell transfer from the GC model, as lymphocytes in F1 mice are tolerized by antigens from both strains. To test this method, we purified splenic B cells by Magnetic Cell Separation (MACS) (S5A Fig) and transferred 2.5x10^5^ B cells from the GC model into each F1 mouse. Hereafter, GC model and F1 mice are referred to as donor and host respectively in adoptive transfer experiments. Two days after B cell transfer, we determined the frequency of donor B cells in host spleen. Donor B cells from the GC model were CD45.1^-^CD45.2^+^. Because the F1 mice were progenies of 129Sv (CD45.2^+^) and C57BL/6.SJL (CD45.1^+^) mice, host B cells were CD45.1^+^CD45.2^+^. Therefore, CD45.1 status can distinguish donor (CD45.1^-^) from host (CD45.1^+^) B cells. Staining by eOD-GT8 added another marker for donor B cells that expressed GL-VRC01. Based on the two markers (eOD-GT8^+^CD45.1^-^), the frequencies of donor B cells in naïve B compartment (B220^+^IgD^+^) of the recipient mice ranged from 1.5-9.8x10^-5^, with a median value at 2.9x10^-5^ (S5B and S5C Fig).

Next, we tested whether donor B cells could respond to immunization in the adoptively transferred mice. Since antigenic stimulation should amplify donor B cells, we further titrated down the number of transferred B cell to 1x10^5^ per mouse, which is 2.5-fold lower than that in the test above and should result in a donor B cell frequency of around 1/85,000 in recipient. Two days after adoptive transfer, we immunized the recipient mice with CH505.core plus alum adjuvant (Fig 4A) and, two weeks after the prime immunization, we analyzed IgG1^+^ GC B cells (B220^+^CD38^lo^GL7^hi^IgG1^+^) (Fig 4B). The rationale to focus on IgG1^+^ B cells was based on the known preference of alum for stimulating Th2 response, which promotes class switching to IgG1 (52). IgG1^+^ GC B cells were further separated into CD45.1^-^ donor B cells and CD45.1^+^ host B cells. This assay revealed a distinct population of donor B cells in GCs, with a median frequency of around 7% from multiple experiments (Fig 4B and 4C). Compared to the starting frequency of 1/85,000 in naive B cells after adoptive transfer, donor B cells have undergone substantial expansion in GC. According to design (Fig 1), donor GC B cells should turn on IA-VRC01 expression. To validate this point, we sorted single IgG1^+^ donor GC B cells from 3 mice, amplified their HCs and LCs with single-cell RT-PCR and determined their identity by sequencing (S2 Table) and restriction digest assay, as for naive B cells in the proceeding section. Based on analysis of 108 donor GC B cells sorted from 3 mice, all the donor IgG1^+^ GC B cells expressed IA-VRC01HC, and, except for one GL-VRC01LC B cell, all expressed IA-VRC01LC as well (Fig 4D, S2, S3 and S4 Fig). Positive selection of IA-VRC01 GC B cells by the prime immunogen likely contributed to the almost complete switching of GL to IA-VRC01, as deletion efficiency of floxed-cassette by the mini-AID cre was approximately 30%, as determined by the YFP reporter (S1 Fig).

**Fig 4.**
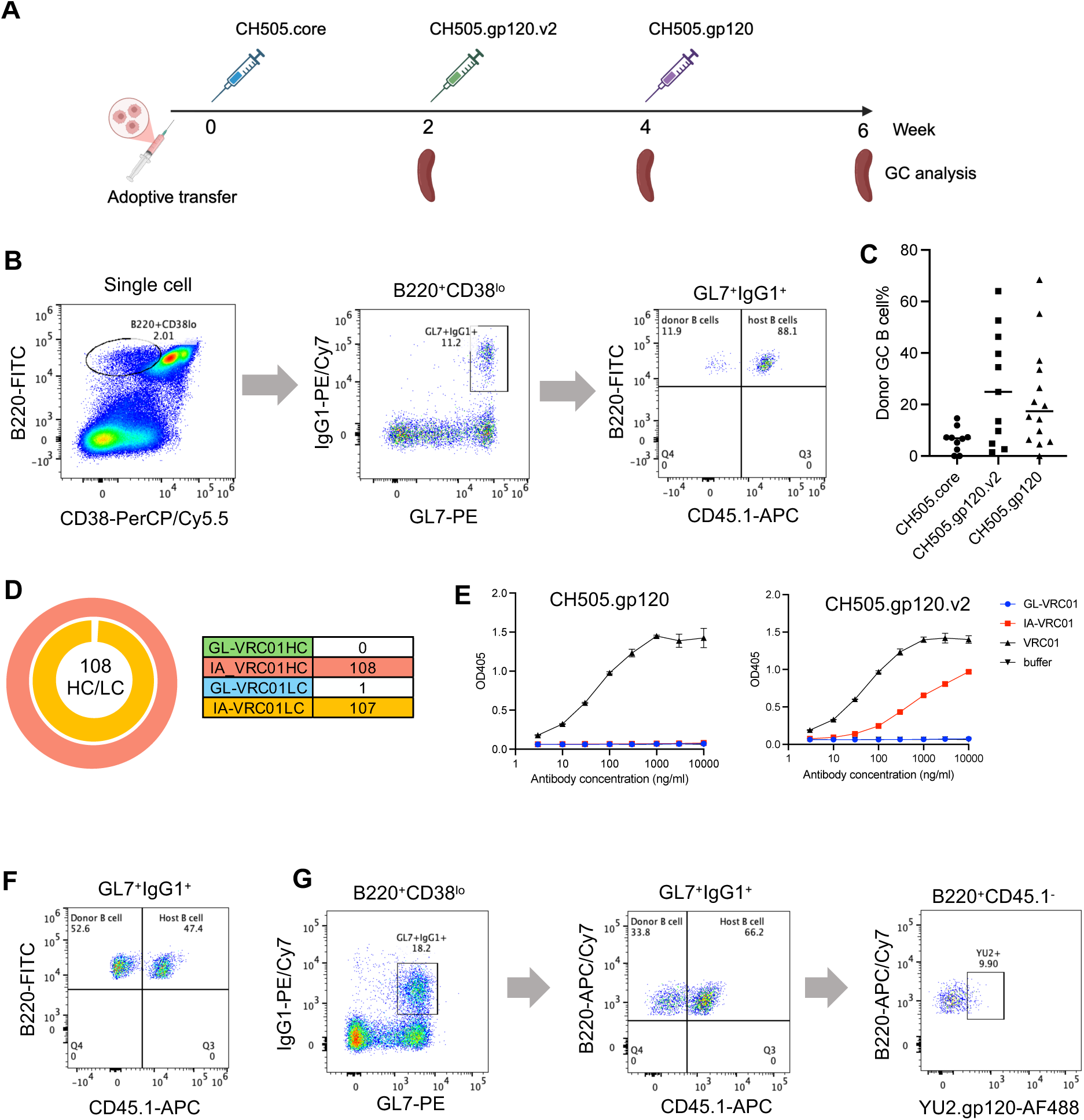
Immunization results of the GC model. (A) Immunization scheme. (B) Flow cytometric detection of donor B cells in GC after prime immunization with CH505.core. Gating scheme: lymphocyte>live cells>single cells>B220^+^CD38^lo^>GL7^+^IgG1^+^>B220^+^CD45.1^-^ (donor GC B cells), B220^+^CD45.1^+^ (host GC B cells). (C) Summary of donor GC B cell frequencies after immunization. The dot plot is based on flow cytometric analysis, as shown in panel B (CH505.core), F (CH505.gp120.v2), G (CH505.gp120). Each dot represents one mouse. The line corresponds to the median. (D) sc-RT-PCR analysis of donor IgG1^+^ GC B cells. The pie chart summarizes the proportion of HC (outer ring) and LC (inner ring) that correspond to GL-VRC01 or IA-VRC01. In total, 108 HC/LC pairs from 3 mice were analyzed. Color codes are shown in the adjacent table, which also lists the number of HC and LC in each category. The shade of the color over the number correlates with HC or LC frequency: 100%, full color; 0%, no color. The identifies of the HC and LC were determined by Sanger sequencing (S2 Fig) or restriction digest (S3 Fig) of the PCR product. The full data are in S4B Fig. (E) ELISA measurement of interactions of GL-VRC01, IA-VRC01 and mature VRC01 antibodies with CH505.gp120.v2 and CH505.gp120. The plots show ELISA titration curves, which are color coded for GL-VRC01, IA-VRC01 and VRC01, as indicated to the right; buffer served as the baseline. Coating antigen is indicated at the top of each plot. The curve corresponds to the average of 3 technical replicates, and the error bar represents standard deviation. (F) Detection of donor GC B cells after the first boost immunization with CH505.gp120.v2. Gating scheme is the same as in panel B. (G) Sorting of donor GC B cells after the second boost immunization with CH505.gp120. Gating scheme is the same as in panel B and F, except for an extra plot at the end: the YU2^+^ population was sorted.

### Maturation of IA-VRC01 by boost immunization

The N276 glycan is a major barrier for VRC01 class antibodies, including the IA-VRC01 in the GC model. To advance VRC01 intermediate beyond this obstacle, boost immunogen should include the N276 glycan to select for accommodating mutations. However, unmodified CH505.gp120, which contains the N276 glycan, showed no detectable interaction with IA-VRC01 by ELISA (Fig 4E, left plot) and probably would not stimulate IA-VRC01 GC B cells. To address this issue, we mutated the glycosylation site at Env position N463 near the CD4 binding site. Relative to the N276 glycan, the N463 glycan is less obstructive for VRC01 class antibodies (47, 53). Nonetheless, mutating the N463 glycosylation site notably improved interaction between CH505.gp120.v2 (CH505.gp120 with N463 glycan deletion) and IA-VRC01 (Fig 4E, right plot). Importantly, the antigen showed an affinity gradient between mature and IA-VRC01, which can select for maturation mutations.

We boosted the mice with CH505.gp120.v2 2 weeks following prime immunization (Fig 4A); the short interval was chosen to keep the donor B cells in GCs for continuous affinity maturation. The first boost immunization further expanded donor GC B cells (Fig 4C and 4F). Following this step, we did a second boost immunization with unmodified CH505.gp120 that contained both the N463 and N276 glycosylation sites (Fig 4A). The second boost immunization did not increase the frequencies of donor GC B cells relative to the first boost immunization (Fig 4C and 4G). We sorted donor GC B cells that stained positive for fluorophore-labeled gp120 from the clade B virus, YU2 (Fig 4G). The heterologous probe could select for BCRs that have broad binding activities. HC/LC pairs from sorted B cells were cloned by single-cell RT-PCR; in total, 125 pairs of HC/LC were obtained from 4 mice (S3 Table). Consistent with IA-VRC01 expression in GC B cells, all 125 HC/LC pairs were derived from IA-VRC01 (S2 Fig). We refer to the HC/LC pairs collectively as IA-VRC01.v2, to distinguish them from the prototype IA-VRC01. Relative to GL-VRC01, IA-VRC01 contains 12.5% amino acid substitutions in HC and 5.7% in LC. Boost immunizations added more mutations into IA-VRC01. With IA-VRC01 as reference, the average amino acid mutation frequencies of IA-VRC01.v2 were 5.9% in HC and 3.4% in LC (Fig 5A). In addition to point mutations, 8/125 (6.4%) HCs had 2aa insertion in CDR H3, 16/125 (12.8%) LCs contained 1-2aa deletion in CDR L1, and 6/125 (4.8%) HC/LC pairs contained both CDR H3 insertion and CDR L1 deletion (Fig 5B). The distributions of point mutations and indels were not random (Fig 5C and 5D), likely reflecting antigenic selection and target preference of AID.

**Fig 5.**
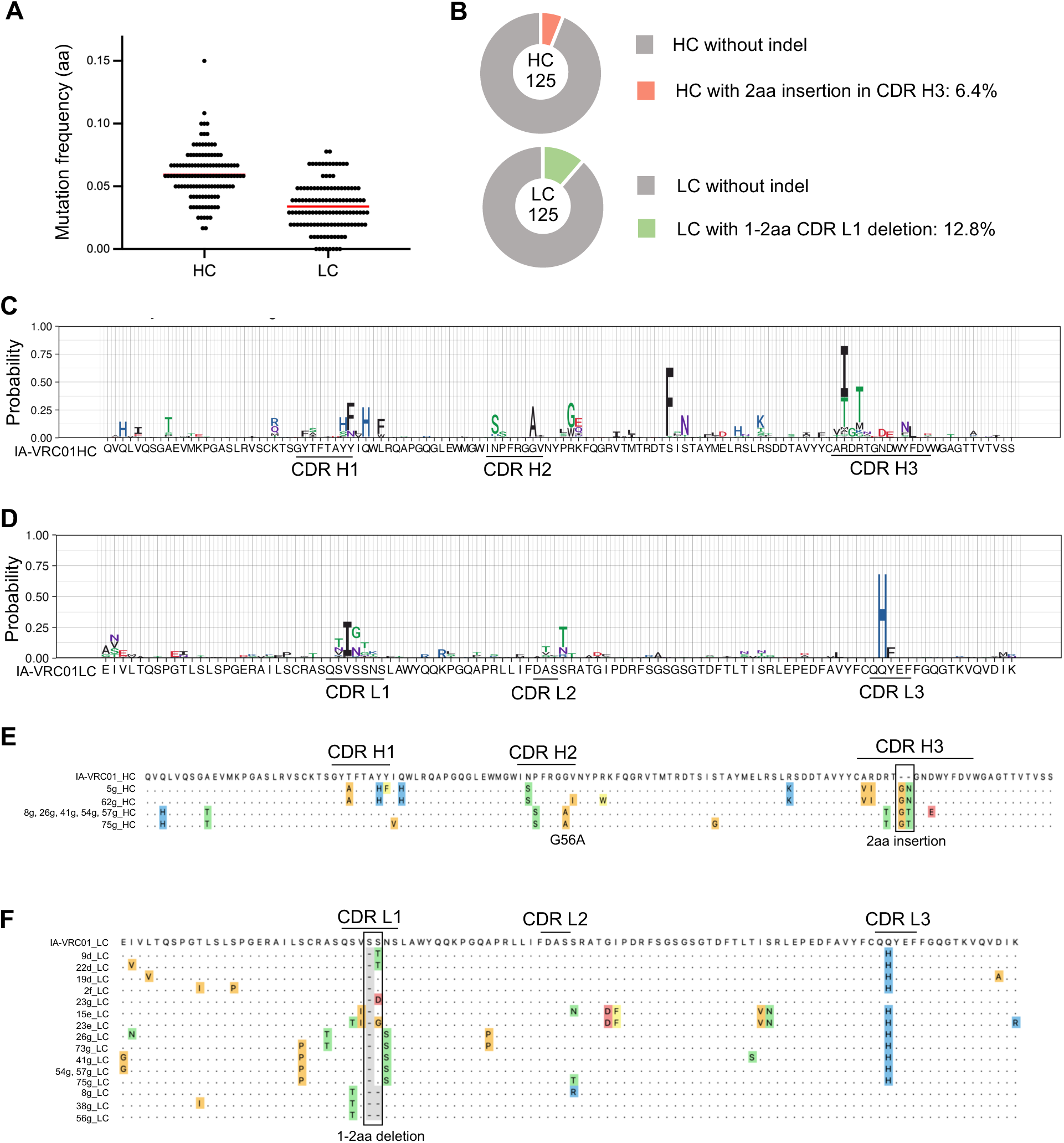
Mutation analysis of IA-VRC01 after boost immunization. (A) Point mutation frequencies of HC and LC. The dot plot shows the amino acid (aa) mutation frequencies of 125 pairs of HC and LC that were amplified by sc-RT-PCR from sorted donor GC B cells of 4 immunized mice (Fig 4G). Each dot represents one HC or LC. The red line represents the median mutation frequency. (B) Indel frequencies of HC and LC. The pie chart summaries the frequencies of CDR H3 insertion and CDR L1 deletion in 125 pairs of HC and LC. (C) Mutation pattern of HC. The logo plot shows amino acid point mutations at each position of the HC variable region, with the original IA-VRC01HC as reference, shown below. Amino acids are represented by standard single letters; the color codes correlate with side chain chemistry. The height of the letter correlates with the frequency of the mutation. The locations of CDR H1, H2 and H3 are marked underneath the logo plot. (D) Mutation pattern of LC. (E) Insertion in CDR H3. HCs with insertion were aligned to the IA-VRC01HC. Each antibody is identified by a number and letter, shown to the left of the alignment; the letter refers to the mouse from which the antibody was isolated (S6 Fig). Sequence identity is indicated by “.“; mutated amino acid residues are shown and color shaded. The VRC01 class mutation, G56A, and the 2aa CDR H3 insertion are marked on the alignment. (F) Deletion in CDR L1. LCs with CDR L1 deletion (“-“ shaded in grey) were aligned to the IA-VRC01LC.

Fifteen mutations in the hV_H_1-2 segment have been defined as VRC01-class mutations, which are characteristic of VRC01 class bnAbs (27). The original IA-VRC01 contains 7 of the 15 VRC01 class mutations in the hV_H_1-2 segment (S6 Fig). After boost immunization, 34/125 (27.2%) IA-VRC01.v2 antibodies gained another VRC01 class mutation: G56A (Fig 5C and S6 Fig). The G56A mutation lies at the core of CDR H2 interface with the CD4 binding loop and is present in most VRC01 class bnAbs. The LCs of VRC01 bnAbs often contain truncations or point mutations in CDR L1, which can reduce steric clash with the N276 glycan. As mentioned above, 12.8% of IA-VRC01.v2 LCs contained 1-2aa deletions in CDR L1 (Fig 5B). Some recurrent mutations in IA-VRC01.v2 antibodies have not been observed before in VRC01 class antibodies (Fig 5B, 5C and 5D). Among these, the 2aa insertion in CDR H3 was notable. An insertion of “GT” residues was present in 6/125 CDR H3s. 5 of the 6 HCs (IA-VRC01.v2_8g, 26g, 41g, 54g, 57g; hereafter, specific IA-VRC01.v2 antibody is referred to by its identification numbrer, for example “8g”) with the “GT” insertion shared the same mutation pattern throughout the HC variable region, and 1 of the 6 (75g) had 2 more mutations (Fig 5E). The 54g and 57g HCs were paired with the same LC; 8g, 26g, 41g HCs were paired with different LCs (Fig 5F). Intriguingly, all these LCs had CDR L1 truncations, including a 2aa deletion in 8g LC. Strong antigenic selection was likely responsible for the coincidence of deletion and insertion, which are infrequent during somatic hypermutation. Two of 125 CDR H3s (5g, 62g) contained an insertion of “GN” residues at the same position of the “GT” insertion (Fig 5E), but the LC partners for these two HCs did not contain CDR L1 deletion (Fig 5F). The CDR H3 insertions were found only in one immunized mouse (S6 Fig) and were likely a rare event. Nonetheless, the CDR H3 insertion appeared to have precipitated clonal expansion, presumably owing to its favorable effects on antigen binding.

We expressed 31 HC/LC pairs as recombinant antibodies for functional analysis. These antibodies had acquired new VRC01-class mutations, indels or other recurrent mutations. We used ELISA to test the interaction of these antibodies with eight Env trimers, which consisted of soluble ectodomains (S3 Table). To quantify binding activity, we calculated the Area Under Curve (AUC) of the ELISA titration curve of each antibody. The AUC ratio of test antibody/VRC01 represented its binding activity, relative to VRC01 as reference. In this assay, the GL-VRC01 showed no detectable binding to any of the Envs (Fig 6A). IA-VRC01 exhibited moderate affinity for 426c Env, as it was originally induced by 426c derived immunogens; its binding to the other Envs was weak (JRFL, YU.2) or undetectable (CH505, CH848, QJX41594.1, AZ172138.1 and ZM106.9) (Fig 6A). After boost immunization, most IA-VRC01.v2 antibodies showed broader and stronger binding to the Env proteins (Fig 6A). For quantitative comparison, we summed up the relative binding activities of each antibody for all eight Envs and defined the value as total binding activity (Fig 6B). Based on this value, 30 of 31 of IA-VRC01.v2 antibodies showed higher Env binding activities than the prototype IA-VRC01. The top four antibodies (8g, 41g, 57g and 26g) bound 7/8 Envs at comparable levels to mature VRC01 (Figure 6A). These four antibodies shared the same HC, which included the “GT” insertion in CDR H3 and the VRC01 class mutation, G56A (Fig 5C). The 2aa insertion was functional, since its removal (8g.1) reduced Env binding activities (compare 8g and 8g.1, marked by *, Fig 6A and 6B). In previous studies, insertions have been found in CDR H1, CDR H3 and framework region of VRC01 class antibodies (13, 21, 54–56). Each of these insertions, including the present 2aa insertion, is unique, likely because insertions are infrequent and stochastic during somatic hypermutation.

**Fig 6.**
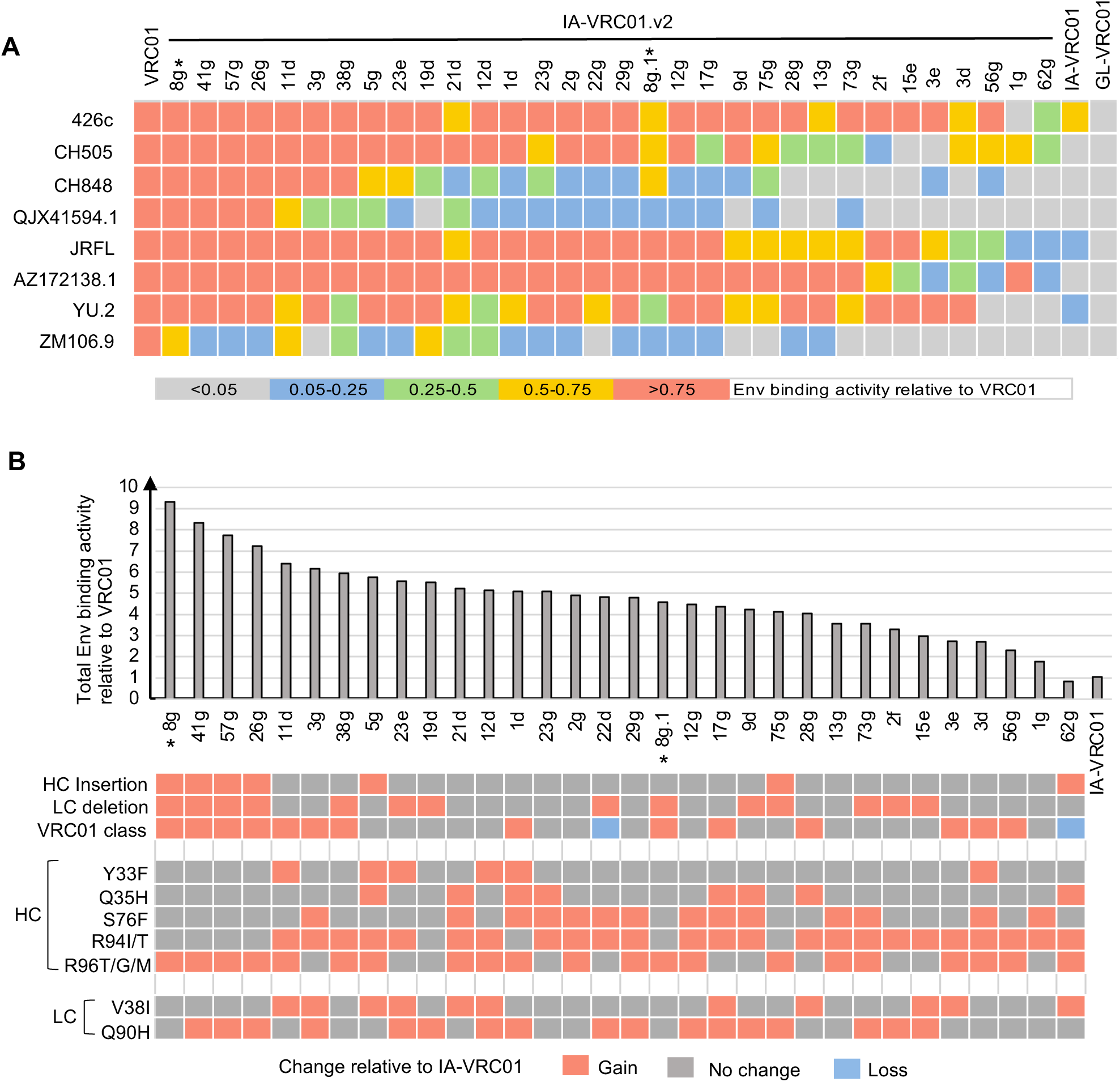
Env binding activities of IA-VRC01 antibodies after boost immunization (IA-VRC01.v2). (A) Binding activities of IA-VRC01.v2 antibodies to Env trimers. The plot summarizes the binding activities of 32 IA-VRC01.v2 antibodies (antibody identification numbers listed on top) to 8 Env trimers (viral strain name listed to the left); for comparison, the plot also includes mature VRC01, the original IA-VRC01 and GL-VRC01 antibodies. 8g.1 was derived from 8g by removing the 2aa CDR H3 insertion; 8g and 8g.1 are marked with * below the plot. Binding activity was measured with ELISA, where Env trimer was the coating antigen. AUC for each antibody titration curve was calculated. The AUC ratio of each antibody/VRC01 represents the relative binding activity and is color-coded on the plot; color code legend is below the plot. The data are average of 2 technical replicates. (B) Correlation of Env binding activity with mutations. The bar graph shows the sum of the binding activities to the 8 Env proteins for each antibody, as measured in panel A. The antibodies are arranged according to total binding activities. The status of mutation is indicated below the bar graph: HC insertion, 2aa CDR H3 insertion; LC deletion, 1-2aa CDR L1 deletion; VRC01 mutation, VRC01 class mutations in hV_H_1-2; HC and LC mutations, recurrent non-VRC01 class mutations in HC and LC. Increase or decrease of mutations are relative to IA-VRC01.

The top four IA-VRC01.v2 antibodies also contained a CDR L1 deletion and the G56A VRC01 class mutation. Antibody 8g contained a deletion of 2aa (S29-S30) in CDR L1. The LCs of antibodies 26g, 41g, 57g shared a deletion of S29 and N31S mutation in CDR L1, but they contained different mutations elsewhere (Fig 5F). Antibody 54g was not included in the ELISA experiment, because it was identical to 57g. The related antibodies (8g, 26g, 41g, 54g, and 57g) may be members of a clonal family that was expanded by virtue of their strong antigen-binding. None of the other antibodies (11d-62g, Fig 6B) had all three mutation events: CDR H3 insertion, CDR L1 deletion and G56A substitution, and these antibodies had lower binding activities. These correlations suggested that the combination of the three mutation events was required for strong Env binding activity. Some recurrent mutations in IA-VRC01.v2 were not VRC01-class, and we did not find any obvious correlation between these mutations and Env binding activity (Fig 6B).

Based on the ELISA results, we picked the top three antibodies (8g, 26g, 41g) for neutralization assays; we did not test 57g, which differed from 41g by only 1 mutation and showed similar binding activities as 41g. The original IA-VRC01 and mature VRC01 antibodies were included in these assays for comparison. The antibodies were tested on a global multi-clade panel of 119 strains of HIV-1 Env pseudoviruses; most of the pseudoviruses have tier 2 phenotypes, which are typical of circulating strains and associated with closed Env configuration (S7 Fig). At 50μg/ml IC50 cut-off, the three IA-VRC01.v2 antibodies neutralized 31.1% (8g), 16.8% (26g) and 27.7% (41g) of the viruses (Fig 7A and S7 Fig). For comparison, the prototype IA-VRC01 had neutralization breadth of 1.7%, and the mature VRC01 neutralized 89.9% of the panel. Relative to the original IA-VRC01, the three IA-VRC01.v2 antibodies improved their neutralization breadth by 18.5-fold (8g), 10-fold (26g), and 16.5-fold (41g), respectively. The geometric means of IC50 were 3.08μg/ml (8g), 7.70μg/ml (26g), and 4.40μg/ml (41g), whereas the geometric mean of IC50 for VRC01 was 0.46μg/ml; based on IC50, the potency of the IA-VRC01.v2 antibodies were still low. Most of the viral isolates that were sensitive to the IA-VRC01.v2 antibodies contain the N276 glycosylation site: 33 of 37 for 8g; 16 of 20 for 26g; and 29 of 33 for 41g. The two viruses that were sensitive to the prototype IA-VRC01 lack the N276 glycosylation site, confirming that these antibody could not accommodate the N276 glycan, as in previous analyses (16). The overall neutralization results showed that the boost immunization enabled some antibodies to overcome the N276 glycan barrier and acquire heterologous neutralization breadth. VRC01 class antibodies with heterologous neutralizing activities have also been elicited in a previous mouse immunization experiment (20). In that experiment, the most advanced VRC01 antibodies could neutralize more than 50% of a 208-virus panel, with IC50 in the μg/ml range. The previous immunization experiment started from a germline precursor and involved multiple immunization steps. It remains to be determined how the boost immunization scheme, devised in the GC model, would work in the context of a complete scheme of prime/boost sequential immunization.

**Fig 7.**
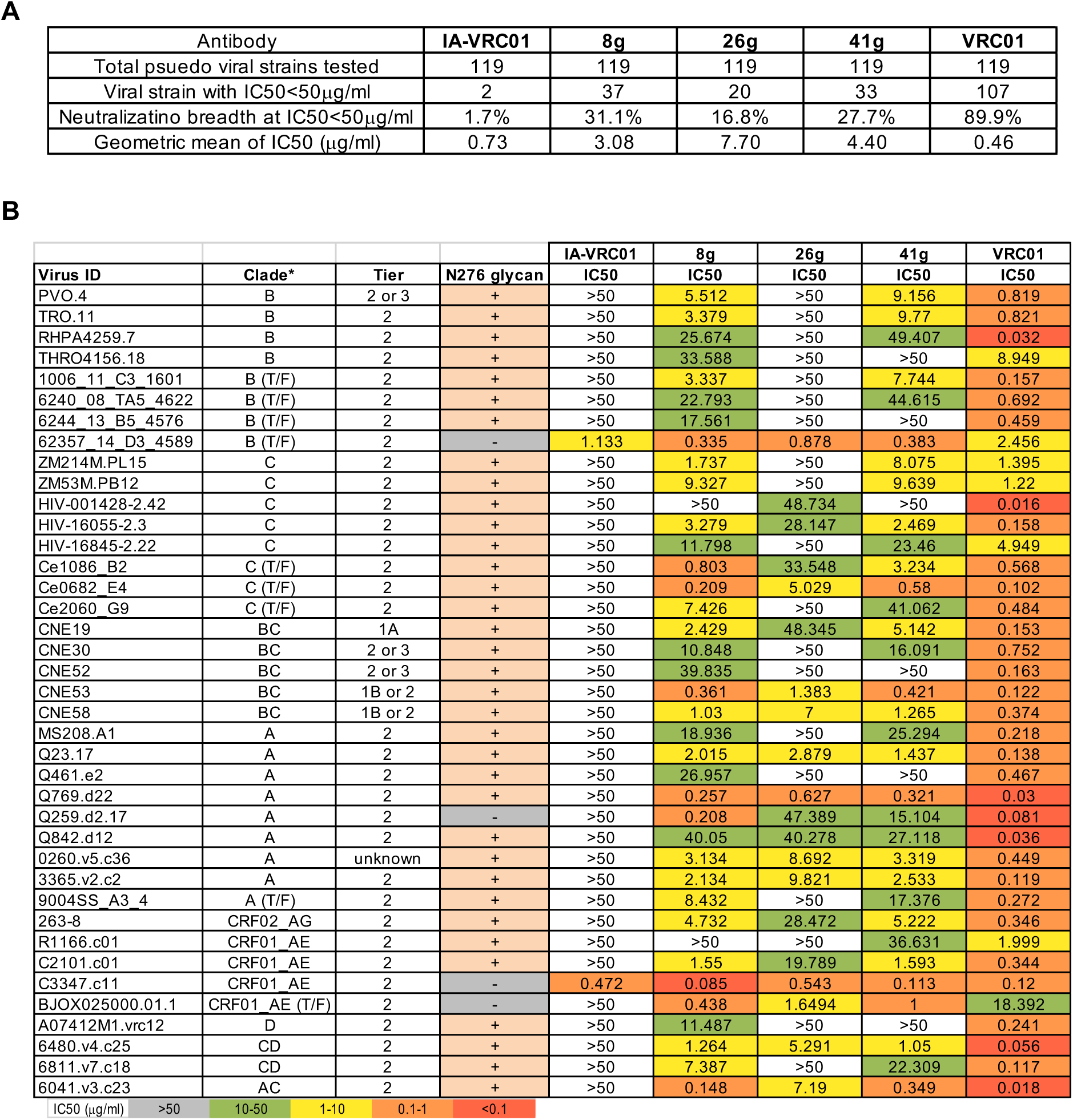
Neutralization activities of IA-VRC01 antibodies after boost immunization. (A) Summary of neutralization results against a global panel of 119 strains of HIV-1 Env pseudovirus. The assay included mature VRC01 and the original IA-VRC01 for comparison. The full neutralization data is in S7 Fig. (B) Details of the sensitive viruses for 8g, 26g and 41g. The table includes only the viruses that are neutralized by 8g, 26g and 41g. Under clade designation, T/F indicates transmitted/founder isolates.

## Discussion

In this work, we generated and characterized a mouse model that was designed to test boost immunization conditions for maturing bnAb intermediates. The present mouse model relied on the GL-VRC01 to support B cell development up to naïve B cell stage, thereby allowing IA-VRC01 expression to bypass tolerance control checkpoints. The strategy would not work in cases where progenitor B cells expressing bnAb precursor are blocked by developmental checkpoints. The problem could be common for bnAbs with long CDR H3s, as seen in multiple knock-in mouse models (4–7). In such cases, one potential solution is to use an innocuous antibody to drive B cell development, as we previously did for conditional expression of CAP256-VRC26 UCA in naive B cells (7). If a “driver” BCR is expressed on naïve B cells, its cognate antigen should be the prime immunogen to induce the bnAb intermediate in GC. However, if the driver BCR is unrelated to the bnAb intermediate, the driver antigen will not engage GC B cells that express bnAb intermediate, and without antigenic support, the nascent GC B cells may not survive to the subsequent boost immunization. One potential way to address this issue is to include the boost immunogen in the priming step, together with the antigen for driver BCR. This way, the boost immunogen will be available for the nascent bnAb intermediate GC B cells.

As a test, we used the GC conditional expression mouse model to advance a VRC01 intermediate beyond the N276 glycan barrier. Because this step represents a big jump in affinity maturation, we used a bridge immunogen. The key feature of this immunogen was removal of the N463 glycan but retention of the N276 glycan. Deletion of the N463 glycan enabled sufficient interaction of the immunogen with the IA-VRC01, whereas the N276 glycan exerted selective pressure to overcome the hurdle. It is unknown whether the strategy will be effective for other VRC01 intermediates that are blocked by the N276 glycan. The effect of N463 glycan deletion may vary for different Envs and VRC01 intermediates. Truncation of variable loops or deletion of other glycans near the CD4 binding site are possible alternatives. Although these modifications alone are not sufficient for germline targeting of precursors, they may be adequate for engaging VRC01 intermediates that have improved affinity for Env. Furthermore, since VRC01 intermediate B cells should have undergone clonal expansion within GC, their frequency in the GC microenvironment could be substantially higher than those of germline precursors in total naïve repertoire, and the relative abundance of VRC01 intermediate B cells could lower the affinity thresholds for boost immunogen.

The main purpose of the GC model is to facilitate boost immunogen design that can be integrated into a complete prime and boost immunization scheme. To serve this purpose, the antibody expressed in the GC model must be representative of the intermediates that arose in sequential immunization. Making the right choice could be challenging, as affinity maturation intermediates in immunization are usually diverse. The dilemma is not unique to the GC model approach, given that all immunogen designs are based on select antibody targets, which cannot encompass all possible variations. Testing more intermediates in GC model would increase the chance of success, but the number needs to practical, as testing each intermediate will entail the generation of a mouse line. Stalled intermediates at major maturation blocks are potentially good choices, because similar intermediates may accumulate at this stage in a reproducible manner. Such stalled intermediates could be identified in sequential immunization experiments in animal models, as in the present study or in human clinical trials. Ultimately, the efficacy of boost immunogen, devised in the GC model, needs to be tested and refined experimentally as part of a prime and boost immunization scheme. Such experiments can be done in mouse models that express bnAb precursors, and many such mouse models are available, including six new models reported in a companion manuscript. The two types of mouse models, GC model expressing bnAb intermediate and rearranging model with diverse bnAb precursors, could serve as complementary tools for vaccine design.

## Materials and Methods

### Generation of the GC conditional expression mouse model

The GC conditional mouse model contains three components: HC expression cassette, LC expression cassette and mini-AID-cre transgene; the complete sequences of the three components are listed in S1 Table. The three components were integrated via homologous recombination into specific loci of ES cells. The HC and LC expression cassettes replaced the mouse J_H_1-J_H_4 and Jκ1-Jκ5 regions respectively. The mini-AID-cre transgene was integrated downstream of the 3’ CTCF binding elements (CBEs) of the IgH locus. In S1 Table, 100bp of genomic sequences flanking the integration sites were shown to help localize the elements in genome.

The ES cell was derived from F1 mouse of 129/Sv and C57BL/6 mouse strains. The IgH loci from the two mouse strains are distinguishable by sequence polymorphisms. Because the homology arms for the integration construct of the mini-AID-cre transgene were derived from the genomic DNA of the 129/Sv strain, homologous recombination took place on the 129/Sv IgH allele. We also used homology arms from the 129/Sv strain to integrate the HC expression cassette on the 129/Sv IgH allele. The linkage of the HC expression cassette and mini-AID-cre transgene simplified mouse breeding to reconstitute the GC model. The LC expression cassette was integrated into the Jκ locus on in a separate ES clone.

Homology-mediated integration was done by transfecting targeting construct into ES cells. In the targeting construct, HC, LC expression cassette or mini-AID-cre transgene were flanked on either side with homology arms for the genomic integration site. The construct contained a Neomycin resistance gene as positive selection marker for stable integration and a Diphtheria Toxin Chain A gene provided negative selection against random integration. To stimulate homologous integration, two guide-RNA/CRISPR expression constructs were designed to cleave the genomic target. The targeting construct and guide-RNA/CRISPR constructs were co-transfected into ES cells. Stable clones were selected with G418 and screened for correct integration by Southern blotting. After the correct clones have been identified, the Neomycin resistance gene was deleted via flanking loxP sites (HC and LC expression cassette) or FRT sites (mini-AID-cre transgene) by transient expression of Cre or Flp recombinase in the ES clones. The step was necessary to avoid interference of local transcription by the Neomycin resistance gene. The deletion was verified with Southern blotting. Finally, the karyotypes of the correct ES clones were analyzed, and clones with normal 40 chromosomes were used for injection.

ES clones were injected into blastocysts that were isolated from Rag2 deficient mice (57). Since Rag2 is essential for V(D)J recombination, the blastocysts themselves cannot produce B and T cells. As a result, all the B and T cells in the resultant chimeric mice originated from the injected Rag2 sufficient ES cells. This Rag deficient blastocyst complementation method allowed us to verify the phenotypes of the various genetic elements in the chimeric stage. After verification, the chimeric mice were crossed with C57BL/6 mice for germline transmission. With this procedure, we generated two mouse lines: HC/mini-AID-cre mice and LC mice. Cross of the two mouse lines generated the complete GC conditional mouse model, which is heterozygous for both the HC/mini-AID-cre allele and LC allele. The GC model was maintained in specific pathogen free facility at Boston Children’s Hospital. Genotyping was done with tail DNA. All the mouse work in this study was approved by the Institutional Animal Care and Use Committee of Boston Children’s Hospital (protocol #00002005).

### Flow cytometry analysis of splenic B cells

Splenocytes were disaggregated into single cell suspensions, stained with fluorophore conjugated antibodies and analyzed on an Attune NxT flow cytometer. FACS plots were generated with FlowJo 10.7.1. The plots were displayed in pseudo color and axis was biexponential. All the antibodies used for the flow cytometry experiments are listed in S4 Table. Following are outlines of the plots and gating schemes in the format: x/y axis (gate). All the FACS analyses were preceded by the following plots and gates: FSC/SSC (lymphocyte) > Sytox blue/FSC (live cells, sytox blue^-^) > FSC-A/FSC-H (single cells) > downstream plots. 1) total B and T cells: CD3/B220 (B cells, B220^+^CD3^-^; T cells, B220^-^CD3^+^). 2) IgM/IgD profile: B220 histogram (B cells, B220^+^) > IgD/IgM (naive B cells: IgM^+^IgD^hi^). 3) Igκ/Igλ distribution: B220 histogram (B cells, B220^+^) > Igκ/Igλ (Igκ^+^ B cells: Igκ^+^Igλ^-^; Igλ^+^ B cells: Igκ^-^Igλ^+^). 4) Follicular and marginal zone B cells: CD93/B220 (mature B cells, B220^+^CD93^lo^) > CD21/CD23 (follicular B cells, CD21^lo^CD23^hi^); marginal zone B cells, CD21^hi^CD23^lo^). 5) VRC01 class B cells: B220 histogram (B cells, B220^+^) > eOD-GT8, DeOD-GT8, CH505.core or DCH505.core histogram (VRC01 class B cells, eOD-GT8^+^/ΔeOD-GT8^-^ or CH505.core^+^/ΔCH505.core^-^). 6) Donor naive B cells in adoptively transferred mice: IgD/B220 (naive B cell: B220^+^IgD^hi^) > CD45.1/eOD-GT8 (donor B cells, CD45.1^-^eOD-GT8^+^). 7) Donor GC B cells in adoptively transferred mice: CD38/B220 (B220^+^CD38^lo^) > GL7/IgG1 (IgG1^+^ GC B cells, GL7^+^IgG1^+^) > CD45.1/B220 (donor GC B cells, CD45.1^-^B220^+^; host GC B cells, CD45.1^+^B220^+^) > YU2.gp120/B220 (YU.2^hi^B220^+^). 8) YFP expression in naive B cells with mini-AID-cre: IgD/B220 (naive B cells: B220^+^IgD^hi^) > YFP histogram. 9) YFP expression in GC B cells with mini-AID-cre: CD38/B220 (B220^+^CD38^lo^) > GL7/CD95 (GC B cells, GL7^+^CD95^+^) > YFP histogram.

### Adoptive transfer and immunization

Adoptive transfer utilized F1 mice as recipient of transferred B cells from the GC model. The F1 mice were progenies of cross between 129SVE (Taconic) and B6.SJL (Taconic) mice. Donor B cells were purified from the spleen of the GC model. The purification employed CD43 microbeads (Miltenyi Biotech), which bind to non-B cells. CD43-labeled cells were depleted by passing through LD column (Miltenyi Biotech), and the flowthrough fraction contained purified B cells (S5A Fig). The purified B cells were diluted to 2.5x10^6^/ml (S5B Fig) or 1x10^6^/ml (Fig 4A) with Phosphate Buffered Saline (PBS), and 0.1ml of cells were transferred to each F1 mouse through intraperitoneal (IP) injection, which can introduce donor B cells into recipient spleen at similar efficiencies as intravenous (IV) injection (58). In our experience, IP injection is technically easier, more reliable, and less stressful for mice than IV injection. Prime immunization began 2 days after adoptive transfer. Immunization was performed through the IP route. Each mouse received 20μg of immunogen together with 50μl of 2% Alhydrogel (InvivoGen) in a total volume of 100μl. Immunizations were separated by 2-week intervals.

### Amplification and analysis of HC/LC pairs from sorted single naive or GC B cells

For isolating single naïve B cells, antibody-stained samples were sorted on a BD FACSAria Cell Sorter. Single cells were sorted into 96-well PCR plates. cDNA synthesis was primed with oligo(dT) primer. To amplify VRC01HC from naïve B cells, the primers anneal to the leader exon of hV_H_1-2 and mCμ. To amplify VRC01LC from naïve B cells, primers anneal to the leader exon of hVκ3-20 and mCκ. It took two rounds of nested PCR to yield detectable cDNA products. S2 Table contains the sequences of the primers and PCR products. To differentiate IA and GL-VRC01HC and LC, the PCR products were digested with RsaI (HC) or NsiI (LC). The restriction sites are indicated on the sequences of the PCR products in S2 Table. The sequences of the PCR products were determined by Sanger sequencing. Their consensus sequence (S2 Fig) was generated by MegAlign Pro of DNASTAR software. Single-cell RT-PCR of GC B cells were done in the same way as for naive B cells, except for the use mCγ1 primer. Mutation analysis of HC/LCs from GC B cells employed MegAlign Pro of DNASTAR. The logo plots of mutations were generated with software: RStudio Version 2025.09.2+418 (2025.09.2+418); package: ggseqlogo and ggplot2.

### Recombinant protein production

eOD-GT8 and DeOD-GT8 proteins were expressed as eOD-GT8-human Fcγ4 fusions, which are dimeric. ΔeOD-GT8 contained D368R and D279K mutations (S5 Table), which abrogate its interaction with VRC01 class antibodies. The recombinant proteins were expressed in Expi293 cells. An Avi tag and 6xHis tag were appended to the C-terminus of hFcγ4 (S3 Table). The 6xHis tag enabled affinity purification of the fusion protein on Ni-column. Protein purification was performed on an AKATA GO FPLC. The recombinant protein was purified from the culture supernatant by Ni-affinity chromatography and gel-filtration. The Avi tag was used to biotinylate the fusion with BirA biotin-protein ligase (Avidity LLC), and biotinylated proteins were conjugated to Streptavidin-fluorophores for use as probes in flow cytometry. This expression, purification and labeling procedures were used for all the proteins in this work. The 426c.core, which was used in ELISA (Fig 3D) was expressed as hFcγ4 fusion. The CH505.core and ΔCH505.core (Fig 3D and 3E) were expressed as mouse Fcγ3 (mFcγ3) fusion, with Avi tag and 6xHis tag at the C-terminus (S5 Table). The choice of mFcγ3 was designed to avoid potential immunogenicity of hFcγ4, as CH505.core was prime immunogen. ΔCH505.core contained D368R mutation in CD4 binding site. CH505.gp120.v2 and CH505.gp120 were also expressed as mFcγ3 for immunization. YU2.gp120, which was used as flow cytometry probe (Fig 4G), was not fused to Fc region. The protein was conjugated directly to AF488 with protein labeling kit from Invitrogen. The Env trimers in Fig 6A were expressed in SOSIP platform with additional stabilizing modifications (sequences shown in S5 Table). To improve cleavage between gp120 and gp41, a human furin expression construct was co-transfected with the Env expression construct. Based on SDS-PAGE analysis, cleavage between gp120 and gp41 was complete for the preparations. The Env trimers were purified via C-terminal His-tag. GL-VRC01, IA-VRC01, mature VRC01 and IA-VRC01.v2 antibodies were expressed in Expi293 cells. The constant regions of the recombinant antibodies were mCγ1 for HC and mCκ for LC. A 6xHis tag was added to the C-terminus of mCγ1, and the recombinant antibodies were purified with Ni-column and gel filtration on FPLC. For ELISA, antigens were coated on plate, and serial dilutions of antibodies were added to the plate. Binding was detected with anti-mouse IgG1-alkaline phosphatase plus its substrate.

### Neutralization assay

The TZM-bl neutralization assay was performed as previously described (59). Briefly, mAbs were serially titrated in duplicate wells using a primary concentration of 50 μg/ml and a 3-fold dilution series. Following addition of HIV-1 Env pseudovirus, plates were incubated for 1 hour at 37° C, followed by addition of TZM.bl cells (1x10^4^/well) in the presence of 11μg/ml DEAE-Dextran (Sigma). Wells containing cells + pseudovirus (without sample) or cells alone acted as positive and negative infection controls, respectively. Following a 48 hour incubation, plates were harvested using Promega Bright-Glo luciferase reagent (Madison, WI) and luminescence detected with a Promega GloMax luminometer. Titers are reported as the concentration of mAb that inhibited 50% or 80% virus infection (IC_50_ and IC_80_ titers, respectively). Virus pseudotyped with the envelope protein of murine leukemia virus (MuLV) was used as negative control. All assays were conducted in a laboratory compliant with Good Clinical Laboratory Practice (GCLP) procedures.

## Acknowledgements

We thank Xuejun Chen, Cheng Cheng and John Mascola at the Vaccine Research Center for suggesting the VRC01 intermediate that was incorporated into the mouse model. We thank Thandi Onami at the Gates Foundation for regular discussions, suggestions and administering the project.

## Author contributions

MT and FWA designed and oversaw the project. MT, JD, HLC, LMT, MET, AW, AY, BW, ID, MS performed experiments. MT, JD, HLC, LMT, MET, BW, ID, MS and FWA analyzed data. MT and FWA wrote the manuscript.

## Data availability statement

All data are included in the manuscript.

## Supporting information

**S1 Fig.**
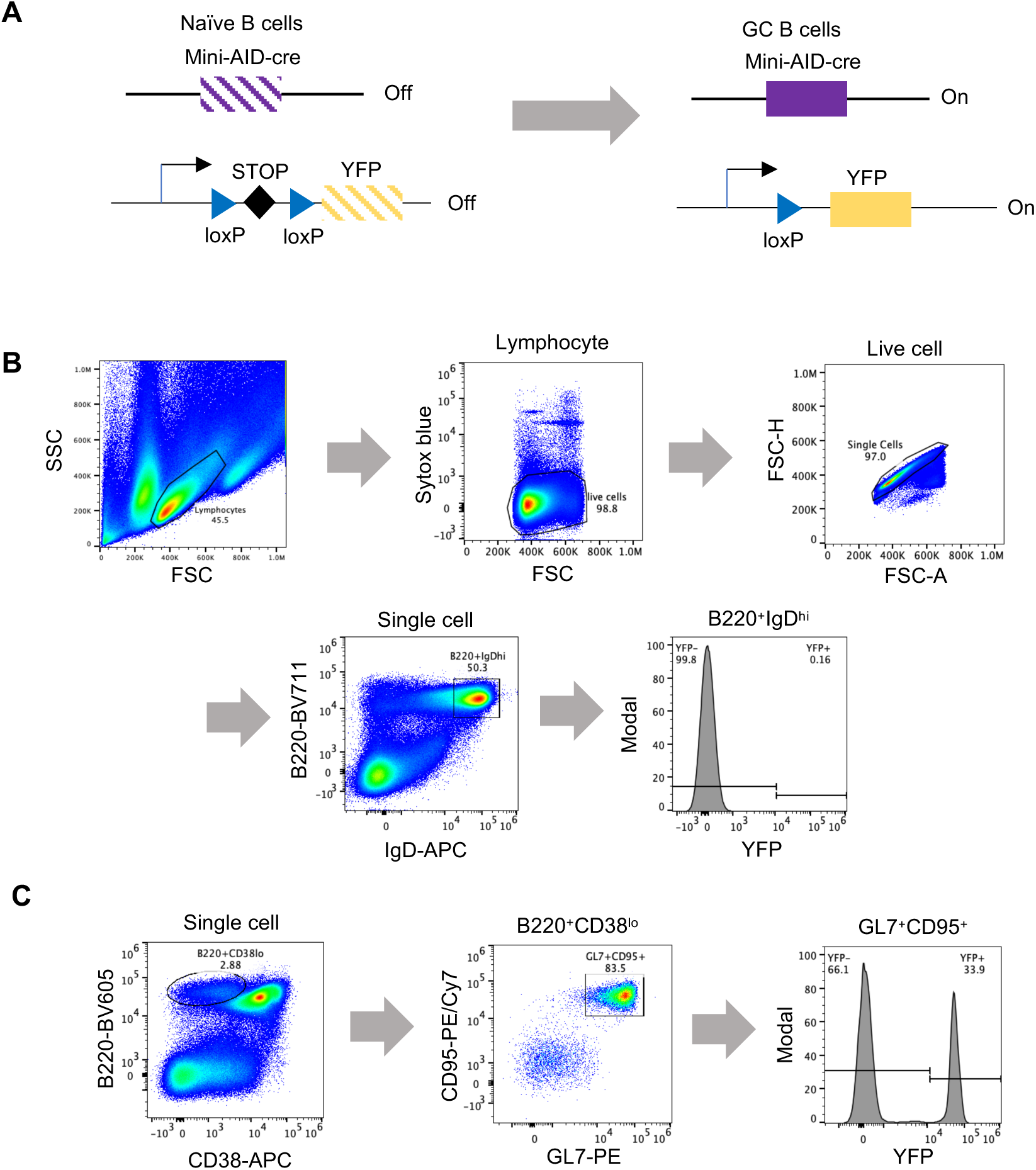
YFP reporter assay for mini-AID-cre. (A) Diagram of the YFP reporter assay. (B) Flow cytometric analysis of YFP expression in naive B cells of unimmunized mouse. Gating scheme: lymphocyte>live cell>single cell>B220^+^IgD^hi^ (naive B cell)>YFP histogram. (C) Flow cytometric analysis of YFP expression in GC B cells after immunization with sheep red blood cells. Gating scheme: lymphocyte>live cell>single cell>B220^+^CD38^lo^>GL7^+^CD95^+^>YFP histogram.

**S2 Fig.**
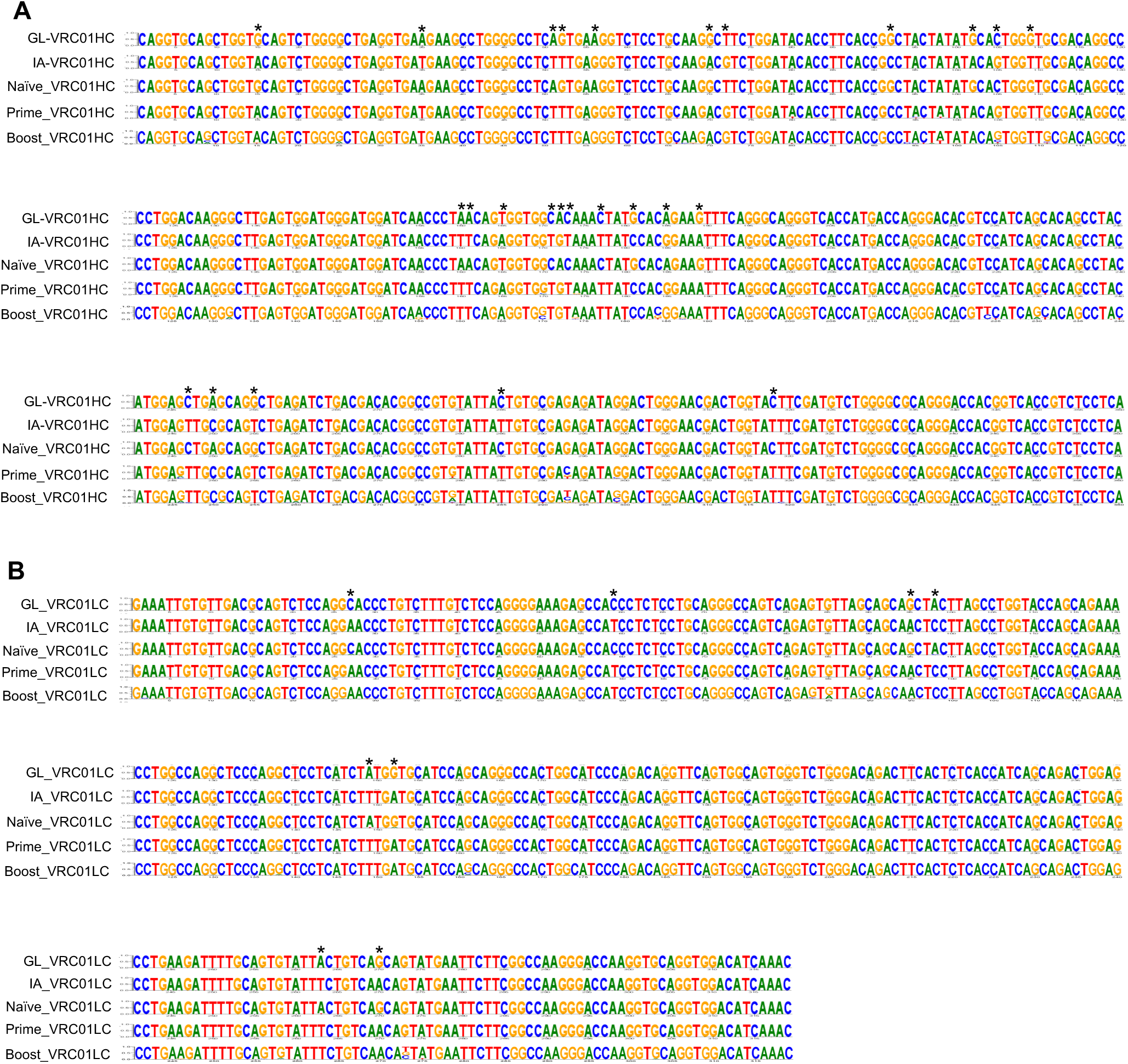
Sequence analysis of VRC01HC and LC from sc-RT-PCR. (A) VRC01HC alignment. The first and second rows are the DNA sequences of the GL-VRC01 and IA-VRC01. Differences between GL-VRC01 and IA-VRC01 are marked with * on top. The sequence of Naive_VRC01HC is the logo plot of all the Sanger sequencing results of sc-RT-PCR products from naive B cells of the GC model (Fig 3C). The height of the residue at each position correlates with its frequency. Prime_VRC01HC is the logo plot of VRC01HC sequences from sorted donor GC B cells after prime immunization of adoptively transferred mice (Fig 4B and 4D). Boost_VRC01HC is the logo plot of VRC01HC sequences from sorted GC B cells after two boost immunizations (Fig 4G). The heterogeneity of residues at some positions reflects somatic hypermutation. (B) VRC01LC alignment.

**S3 Fig.**
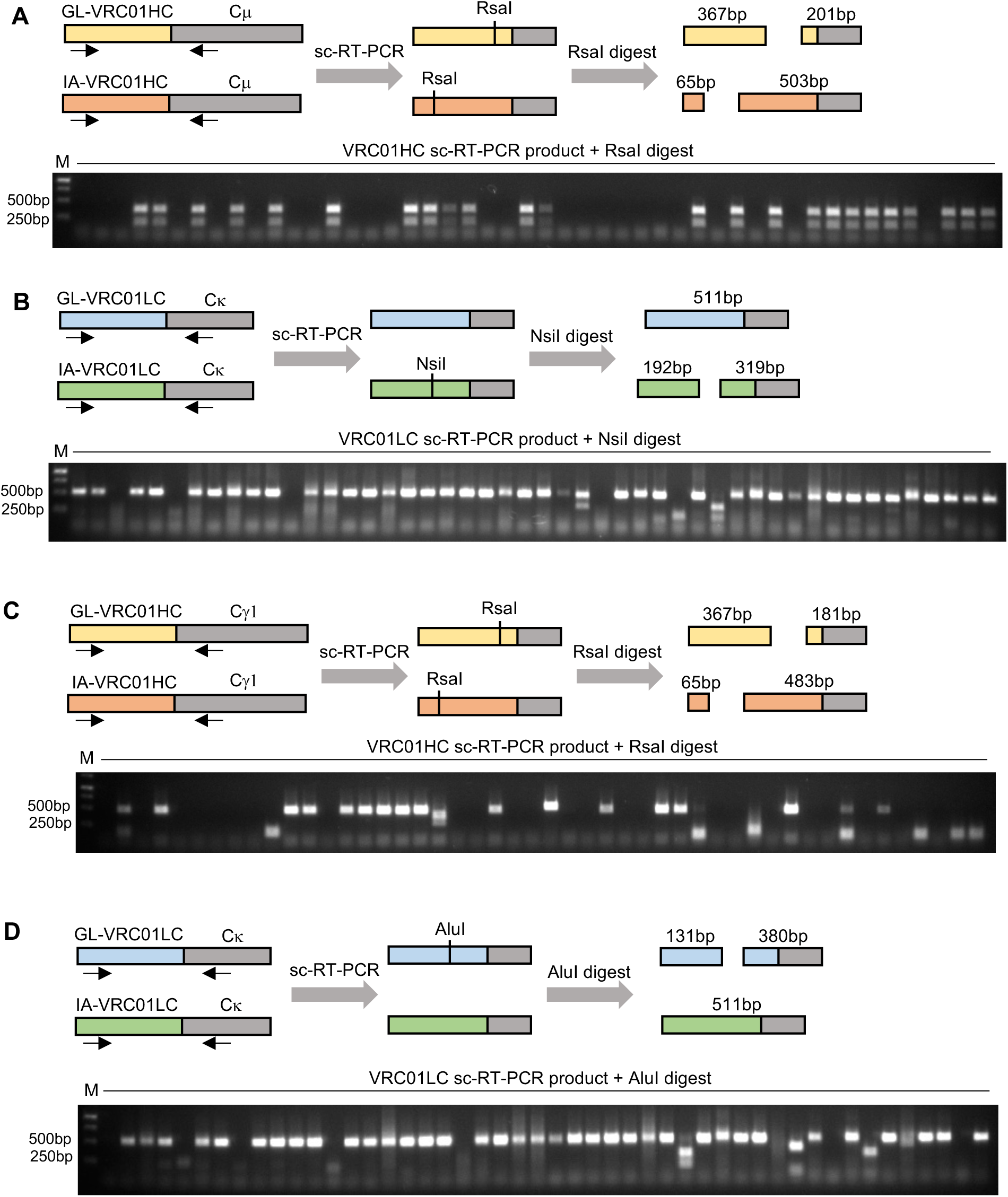
Restriction digest analysis of VRC01HC and LC sc-RT-PCR products. (A) Analysis of VRC01HC from sorted naive B cells of unimmunized GC model (Fig 3C). The diagram shows the position of the PCR primers (arrows) for amplifying VRC01HC cDNA. Due to somatic hypermutation, IA-VRC01HC differs from GL-VRC01HC in RsaI restriction digest pattern. The gel image below is an example of sc-RT-PCR products that have been digested with RsaI. Each lane contains the PCR product of a single cell. Sc-RT-PCR did not work for the cells in empty lanes. (B) Analysis of VRC01LC from sorted naive B cells of unimmunized GC model (Fig 3C). The panel is analogous to panel A. IA-VRC01LC differs from GL-VRC01LC in NsiI restriction digest pattern. (C) Analysis of VRC01HC from donor GC B cells after prime immunization (Fig 4B and 4D). (D) Analysis of VRC01LC from donor GC B cells after prime immunization (Fig 4B and 4D).

**S4 Fig.**
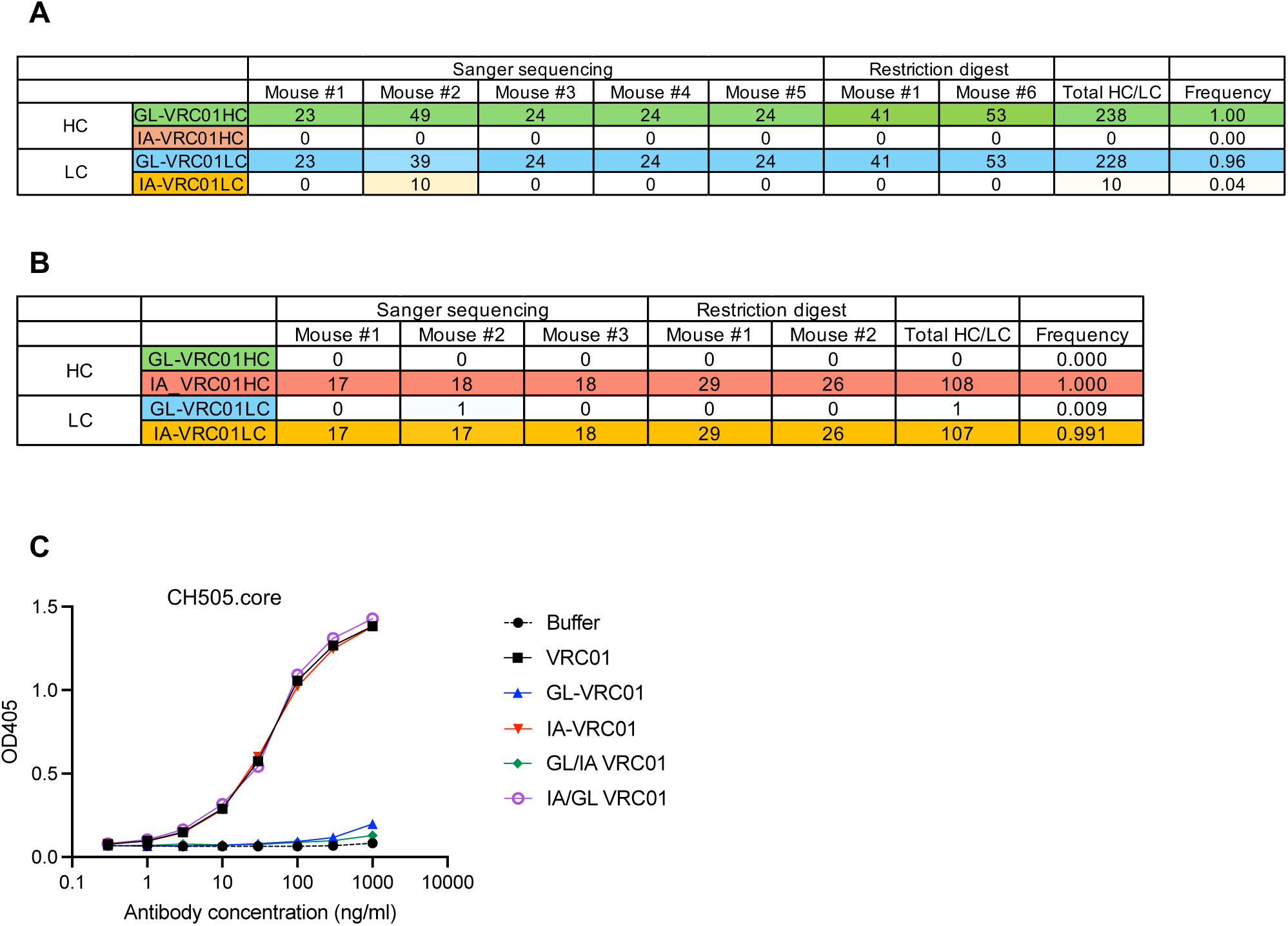
Summary of sc-RT-PCR analysis. (A) sc-RT-PCR results of naive B cells. This table is based on the experiments in Fig 3C, S2 and S3 Fig. The table includes more details than the one in Fig 3C. (B) sc-RT-PCR results of donor GC B cells after prime immunization. This table is based on the experiments in Fig 4B, 4D, S2 and S3 Fig. This table includes more details than the one in Fig 4D. (C) Analysis of the binding activities of hybrid antibodies (GL/IA-VRC01, IA/GL-VRC01) by ELISA.

**S5 Fig.**
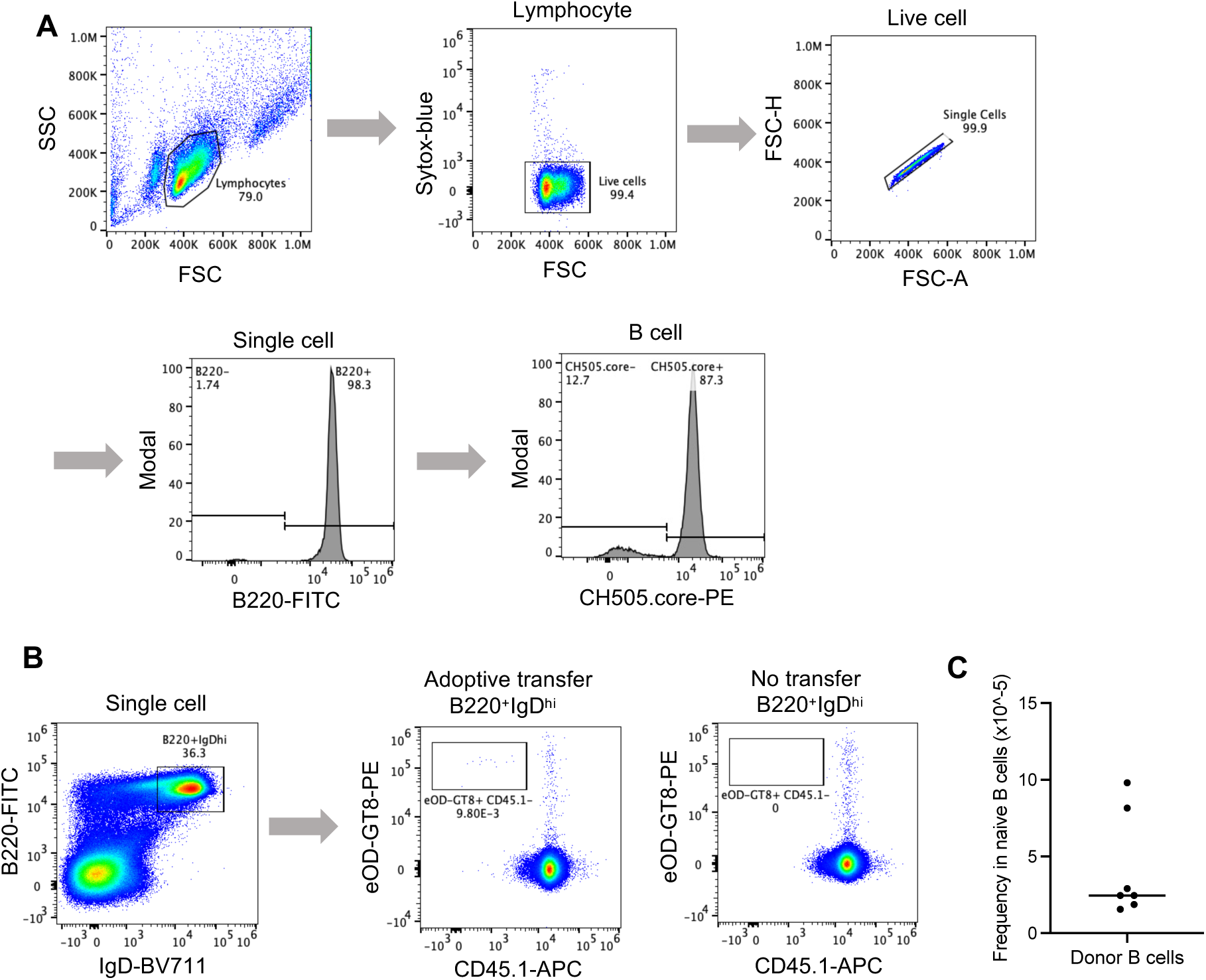
Flow cytometric analysis of adoptive transfer experiment. (A) Flow cytometric analysis of donor splenic B cells from the GC model. Splenic B cells from the GC model were purified with MACS. The FACS plots show the analysis of the purified splenic B cells before adoptive transfer. Gating scheme: lymphocyte>live cell>single cell>B220^+^ B cell>CH505.core^+^ VRC01 B cell. (B) Flow cytometric analysis of donor B cells in the naive B cell compartment of adoptively transferred mice. The analysis was done on splenocytes two days after adoptive transfer. Gating scheme: lymphocyte>live cell>single cell>B220^+^IgD^hi^ (naive B cell)>CD45.1^-^eOD-GT8^+^ (donor B cell). (C) Summary of donor B cell frequency in naive B cell compartment in adoptively transferred mouse. The dot plot is based on FACS analysis as shown in panel B. Each dot corresponds to one adoptively transferred mouse; the line represents the median.

**S6 Fig.**
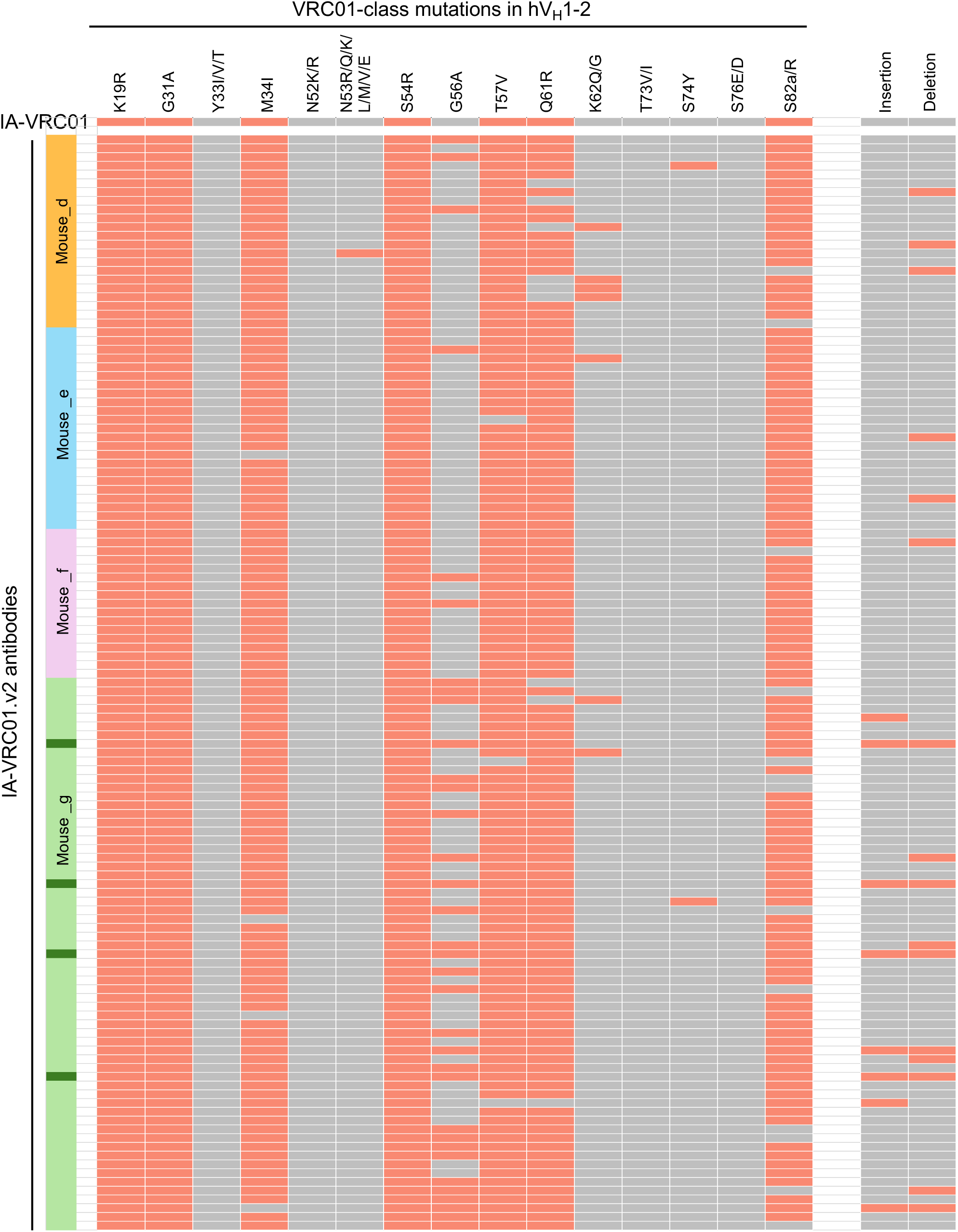
VRC01 class mutations and indels in 125 HC/LC pairs of IA-VRC01 antibodies after boost immunization. VRC01 class mutations in the hV_H_1-2 segment and indels are listed on top; the presence or absence of VRC01 class mutations and indels are represented by red and grey respectively. Each horizontal line represents one antibody. IA-VRC01 is represented by the line at the top. Below are antibodies after boost immunization (IA-VRC01.v2), isolated from 4 mice, and the mouse origin of each antibody is indicated to the left. The mutation status in Fig 6B is based on this plot.

**S7 Fig.**
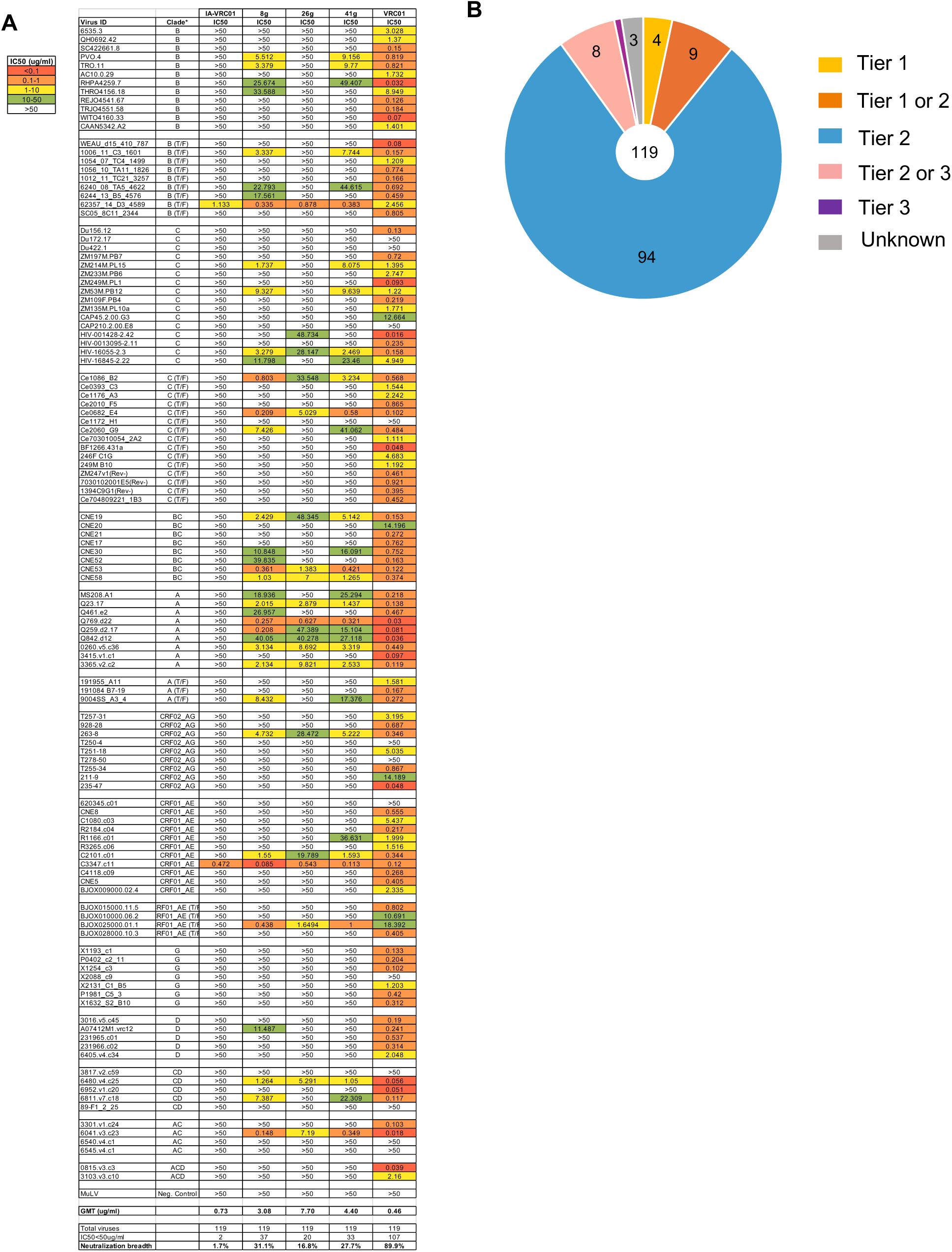
Neutralization results. (A) The table shows the complete neutralization data of the top three IA-VRC01.v2 antibodies (8g, 26g, 41g), the original IA-VRC01 and mature VRC01. Fig 7 is based this data. (B) Tier distribution of the 119 pseudoviruses in the neutralization panel.

**S1 Table. Sequences of the GC model components.**

(A) HC conditional expression cassette. The coding regions for the variable regions of IA-VRC01HC and GL-VRC01LC are in red and blue respectively. 100bp of genomic sequences flanking the expression cassette are shown in italic. (B) LC conditional expression cassette. The coding regions for the variable regions of IA-VRC01LC and GL-VRC01LC are in red and blue respectively. 100bp of genomic sequences flanking the expression cassette are shown in italic. (C) Mini-AID-cre transgene. The coding region for the cre is in red. 100bp of genomic sequences flanking the mini-AID-cre transgene are shown in italic.

**S2 Table. Sequences of the sc-RT-PCR products of GL-VRC01 and IA-VRC01 (Fig 3C, 4D, S2, S3, S4 Fig).**

The primer site is highlighted in red.

**S3 Table. IA-VRC01.v2 antibody sequences.**

The table includes the sequences of the 125 pairs of HC/LC that were characterized in Fig 5-7. The first row contains the sequences of the original IA-VRC01HC and LC.

**S4 Table. Antibodies for flow cytometry**

**S5 Table. Sequences of the recombinant proteins used in this study.**

